# Introduction bias: Imbalance in species introductions may obscure the identification of traits associated with invasiveness

**DOI:** 10.1101/2021.03.22.436397

**Authors:** Estibaliz Palma, Jian Yen, Peter A. Vesk, Monserrat Vilà, Jane A. Catford

## Abstract

The introduction stage is usually overlooked in trait-based studies of invasiveness, implicitly assuming that species introductions are random. However, human activities promote the movement of specific types of species. Thus, species deliberately introduced for distinct purposes (e.g. gardening, forestry) or as contaminants of human commodities (e.g. stowaway) will likely show particular traits. If species with certain traits have been preferentially introduced (i.e. introduction bias), some traits may have been mistakenly linked to species’ invasion abilities due to their influence on introduction probability.

In this work, we propose a theoretical framework with different scenarios of introduction bias. The introduction scenarios are: (1) *Random introduction*, independent from traits; (2) *Biologically biased introduction*, following the worldwide distribution of the trait; and (3) *Human biased introduction*, following a theoretical introduction pathway that favours the introduction of species with high values of the trait. We evaluate how the introduced trait distributions in these scenarios may affect trait distributions in naturalized and invasive species pools under different hypothesized associations between traits and the probabilities of naturalization and invasion. The aim of this work is to identify situations where ignoring introduction bias may lead to spurious correlations being found between species’ traits and species’ ability to become naturalized or invasive.

Our framework strongly points to the need to evaluate the traits of species that have become naturalized or invasive along with the traits of species that have failed to do so in order to unravel any existing introduction bias that may confound the correlation between species’ traits and invasion success. Overlooking a possible introduction bias may lead to the overestimation of the correlation between the trait and the species’ invasion ability, especially in cases when the pool of introduced species shows extreme values of the trait distribution (as compared to a random introduction). Trait-based studies that deserve special attention to avoid undesired effects of introduction bias on their findings are: those that investigate naturalization using only the pool of naturalized species, and those studies that examine invasiveness by comparing invasive species with native species.

## 2 Introduction

Biological invasions result from species moving beyond their natural biogeographic boundaries and establishing self-sustaining populations in the introduced range, as a consequence of human intervention. The set of dispersal, environmental and biotic barriers that need to be overcome for successful invasion has been named the ‘introduction-naturalization-invasion continuum’ (Richardson et al. 2000; Blackburn et al. 2011). Comparative studies aiming to determine invasion drivers, including the effect of functional traits, commonly adopt a stage-based approach, studying the ability of species to successfully reach subsequent stages (e.g. establishment, naturalization, invasion of intact vegetation; Dawson et al. 2009). Trait-based invasion studies commonly focus on the naturalization and/or invasion stages, and attribute species’ unequal success to naturalize or become invasive to differences on their traits (Moravcová et al. 2010; Gallagher et al. 2015). However, functional approaches in invasion science usually overlook the introduction stage (e.g. Hamilton et al. 2005; Chen et al. 2015) due to the scarcity of introduction history records, which may potentially hinder the correct interpretation of observed patterns of naturalization and invasion.

Introduction of species is a key step of the invasion process and the reasons behind the introductions may shape the subsequent stages by limiting the set of species that can become naturalized and, ultimately, invasive. Individuals and species are not randomly transported around the world; rather, humans typically favour the movement of particular functional types (particular species or even particular forms of species) through well-established introduction pathways (Wilson et al. 2009b; Essl et al. 2015). If some types of species are preferentially introduced, it is likely that particular traits or combinations of traits are overrepresented in the pool of introduced species (introduction bias; Kueffer et al. 2013). A potential consequence of the introduction bias is that some traits may have been previously mistakenly linked to invasiveness due to their disproportionate introduction, rather than their influence on invasion success (van Kleunen et al. 2015). Concerns about overlooking the potential species pool are not limited to invasion science: community assembly studies that ignore the whole regional species pool and investigate the processes shaping local biodiversity focusing on the observed local pool alone may also be misleading (Pärtel et al. 2011).

By ignoring the introduction stage, invasiveness studies usually make the implicit assumption that introduction of species is random from a global pool of available species, and therefore independent of plants’ traits and unbiased (i.e. all trait values have the same probability to be introduced). However, a more realistic approach would acknowledge that the introduced species may be biased towards particular taxa or life-history traits (Diez et al. 2009), because they are either preferred for human activities (e.g. gardening, forestry, pastoralism) or accidentally introduced more often, as stowaways or contaminants (Wilson et al. 2009b). Some studies avoid the overestimation of the involvement of traits in invasion due to non-ecological drivers, including the preferential introduction of such traits, by confirming that the traits associated with naturalization/invasion success are also linked (in the opposite way) with species’ failure to become naturalized/invasive (Zenni & Nuñez 2013). Including both successful and unsuccessful species is more common for studies focused on the invasion than the naturalization stage (Moravcová et al. 2010; Gallagher et al. 2015), given that records of all the naturalized species (including invasive and non-invasive) are usually available but records on species that have failed to naturalize after introduction are rare to find (but see Lavoie et al. 2016; Diez et al. 2009). Without formally incorporating any existing introduction bias into the analyses, studies that compare successful and unsuccessful species provide information on traits that are linked to invasiveness conditional on introduction. The extent of introduction bias has rarely been addressed in the literature (but see Diez et al. 2009; Martin et al. 2009; Chrobock et al. 2011), with results from early works suggesting important implications for trait-based invasiveness studies (Kitajima et al. 2006; Martin et al. 2009).

In this work, we aim to identify situations where ignoring introduction bias can lead to spurious correlations being found between species’ traits and their ability to become naturalized or invasive. We first describe different scenarios of introduction bias and how they may influence the pool of species that ultimately become invasive. Then, through simulations based on a simple framework (Figure 1), we explore the cascading effects that introduction bias may have on studies dealing with correlations between traits and species’ naturalization and invasion potential. Finally, we focus on real functional traits widely used across plant invasion studies (specific leaf area, height, seed mass and woodiness) and simulate changes on their distributions along the introduction-naturalization-invasion continuum based on our hypothetical introduction scenarios. Disentangling the influence of introduction bias on invasion patterns and trait-based studies is central to improving inference about how traits promote plant invasion.

**Figure 1.**
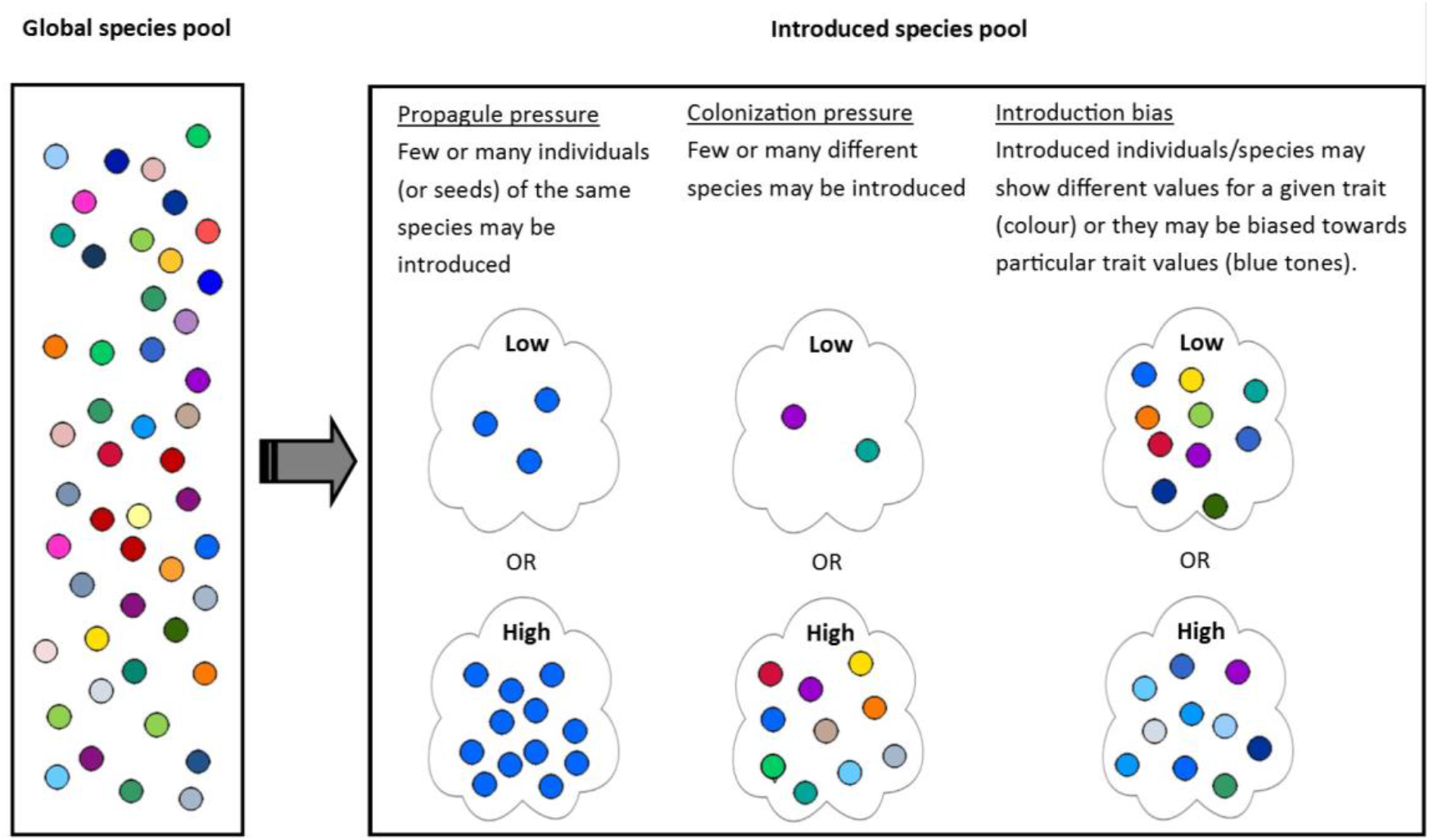
Representation of introduced pools with different degrees of propagule pressure, colonization pressure and introduction bias arising from a hypothetical global species pool.

**Box 1** On the relationship among propagule pressure, colonization pressure, introduction bias and the global pool of species.

Recent research has identified different biases that must be considered to understand biological invasions. These include propagule pressure, colonization pressure and introduction bias (Figure 1). *Propagule pressure* – or introduction effort – is defined as the number of propagules (e.g. individuals, seeds) released into a new environment, and the number and nature of the release events across time and space (Simberloff 2009). Propagule pressure increases the likelihood that a given introduced species overcomes early barriers of invasion and establishes an ongoing population, rather than going extinct due to stochastic demographic, environmental or genetic factors (Lockwood et al. 2005; Duncan 2011). Some authors have further claimed that propagule pressure increased the probability of naturalized species becoming invasive (Essl et al. 2010; but see Dehnen-Schmutz et al. 2007). Although access to reliable, good-quality information on propagule pressure (or its surrogates) is a challenge, invasion ecologists have made great efforts to incorporate it into invasion studies and agree that only species with similar introduction pressure should be directly compared (Mulvaney 2001; Colautti et al. 2006).

*Colonization pressure* is the total number of exotic species released into a single location (Lockwood et al. 2009; Blackburn et al. 2015). At community level, colonization pressure emulates the same pervasive effect that propagule pressure has at the population level; as more species are introduced, more will naturalize due to environmental and biotic suitability by chance alone. For example, Diez et al. (2009) showed that plant families with largest numbers of naturalized species in Australia and New Zealand are those for which the most species have been introduced (e.g. Poaceae, Asteraceae and Fabaceae). Colonization pressure has not received as much attention across the invasion literature as propagule pressure has, but it has been indirectly addressed through comparative studies of native and exotic species richness (communities subject to high human impact are dominated by alien species; Tomasetto et al. 2013).

*Introduction bias* refers to the fact that some species, or types of species, are transported and introduced outside of their native range more often than others. This selection responds to human interest, economic or aesthetic, both direct and indirect. Particular species can be actively chosen because of their value (e.g. agriculture) or they can be accidentally transported because of their close relationship with particular human activities or areas (e.g. introduction by ballast waters). Introduction bias has been studied in the form of introduction pathways (Wilson et al. 2009b) and invasion syndromes (Kueffer et al. 2013). Most of the literature dealing with non-accidental introduction of plants has focused on the horticultural (e.g. Chrobock et al. 2011) and pastoral pathway (e.g. Lonsdale 1994).

Propagule pressure, colonization pressures and introduction bias each play a role in which species from the global pool of species arrive in a new location (Figure 1). One of the main challenges of invasion studies is that introduced species have different propagule pressures and likely come from several donor regions (where they have different relative abundance) at different times (Lockwood et al. 2009), all while human activities shape particular types of introductions.

## 3 Theoretical framework

### 3.1 Invasion stages

Following the concept of the introduction-naturalization-invasion continuum (Richardson et al. 2000; Blackburn et al. 2011), we defined three discrete stages; introduction, naturalization and invasion. Introduced species are those that have overcome major geographical barriers via human activity (Richardson et al. 2000). Naturalized species are a subset of the introduced species able to establish long-term, self-sustaining populations. This definition excludes casual plants (sensu Richardson et al. 2000) due to their short-term persistence and reliance on human inputs. Invasive species are the subset of naturalized species that, having overcome both reproductive and dispersal barriers, establish populations far from the introduction area, with high relative abundance and/or under a wide range of environmental conditions (Catford et al. 2016). We refer to the group of introduced species unable to naturalized as failures (following Diez et al. 2009), and to the naturalized species that never become invasive as non-invasive. In this framework, we follow the terminology proposed by Richardson & Pysek (2011) – i.e. the ‘naturalization-invasion continuum – and limit the use of the term invasion to refer to the last stage of our framework, excluding the previous two stages (i.e. introduction and naturalization).

For simplicity, we summarize invasion processes in only three stages (introduction, naturalization and invasion); however, we acknowledge that these stages may take different names and that more specific stages could be defined. For example, introduction can be broken into uptake/transportation and release/introduction (Heger & Trepl 2003; Lockwood et al. 2005; Blackburn et al. 2011; Lockwood et al. 2013). We are also aware that the declaration of species as invasive can follow other criteria (e.g. impact); however, given our interest in studying the links between invasion and traits, a demographic approach is more appropriate here.

#### 3.1.1 Stage 1: Introduction

The framework integrates the idea that introductions reflect particular introduction motivations (e.g. ornamental, agriculture) and pathways (e.g. mass dispersal and cultivation) (Wilson et al. 2009b), resulting in some functional bias within the pool of introduced species. In this work, we define introduction bias as the preferential introduction of species with particular combinations of traits, regardless of the propagule pressure of each individual species (although they are likely to be positively correlated; Box 1, Panel A1).

We present three possible scenarios of introduction:

##### *Random introduction*: independent of traits

This scenario does not consider the possibility of introduction bias. Introductions are random along the gradient of trait values (i.e. every trait value is introduced with similar frequency). Consequently, introductions cannot be predicted based on species’ traits. This is the implicit assumption routinely made in invasion studies that examine correlations between traits and invasion patterns but do not consider the pool of introduced species (e.g. Hamilton et al. 2005).

##### *Biologically biased introduction*: trait follows the worldwide distribution

This scenario incorporates a form of introduction bias while assuming that species introductions happen randomly. If a particular trait is unrelated to introduction motivations or pathways, the introduced species pool will be a random subset of the global pool of species. As a result, the distribution of that trait in the introduced pool would follow the distribution of that trait in the global species pool. In other words, trait values with higher occurrence worldwide (i.e. more species have them, the species with them are more abundant, or the species with them have larger geographic ranges) will be introduced more often.

##### *Human biased introduction*: trait distribution biased towards preferences of humans or human activities

This scenario reflects that human activities can modify the functional composition of the introduced pool of species in a way that reflects the human motivation behind the introduction (Knapp & Kühn 2012; Essl et al. 2015; Pergl et al. 2017). This applies to accidental (e.g. contaminant, stowaway) as well as deliberate introductions (e.g. agriculture, horticulture). The former introductions will be biased toward species with a strong association with human activities (e.g. weeds of agriculture), showing good dispersal ability to stowaway in human vessels and able to quickly establish after release even in small quantities (Pyšek et al. 2011). For deliberate introductions, different functional biases may be found depending on the reasons behind the introduction. Forestry will promote the introduction of taller species (McGregor et al. 2012); agriculture will promote pastures with high growth rate and good environmental match (Driscoll et al. 2014); and horticulture will promote the introduction of species with low maintenance requirements, ease of propagation and resistance to environmental stress, such as drought or pests (Martin et al. 2009; Drew et al. 2010; van Kleunen et al. 2018b). This scenario represents a simplified, hypothetical human bias towards species that are particularly useful, appealing or associated with human activities. We assume that the bias resulting from human activities and preferences acts on top of the biological bias described in the previous section, meaning that the global distribution of the trait will bound the range and relative availability of trait values over which the human bias operates.

#### 3.1.2 Stage 2: Naturalization

Traits may correlate with the probability that introduced species will successfully naturalize. The framework presents a dichotomy where the trait does or does not affect naturalization. If there is a correlation between the trait and the naturalization success, it is assumed to be positive. That is, the frequency of the successful trait values will increase in the naturalized species pool, regardless of the introduction scenario.

#### 3.1.3 Stage 3: Invasion

As for Stage 2, the framework presents a dichotomy where the trait does or does not affect invasion probability. If there is a correlation between the trait and the probability of become invasive, it is assumed to be positive. That is, the frequency of the successful trait values will increase within the pool of invasive species, regardless of the introduction and naturalization scenarios.

### 3.2 Expectations

The species introduced under the *Random introduction* scenario will show the widest range of trait values while the species introduced under the *Human biased introduction* scenario will have the narrowest range of trait values. The trait distribution will change among stages (from introduction to naturalization to invasion) and between successful and unsuccessful groups within each stage (e.g. naturalized species and failures) only when the trait is correlated with the success of species to become naturalized and/or invasive. The stronger and more coordinated (similar direction) the trait effects on consecutive stages, the larger shift towards narrower distribution of values from introduction to invasion.

The trait distributions of naturalized and invasive species result from the combined relationships of the trait and the probability to reach different stages. Similar distributions of traits for naturalized or invasive species may rise from several combinations of positive, negative and neutral effects of the trait on the probability to be introduced, naturalize and/or become invasive. For example, comparable trait distributions of naturalized species may result as either a combination of a *Biologically biased introduction* and a strong positive effect of the trait on naturalization, or a combination of a *Human biased introduction* towards plants with large values and a neutral or small effect of the trait on naturalization.

## 4 Methods

We first simulated a framework describing the distribution of a theoretical trait for introduced, naturalized and invasive species, based on the three introduction scenarios described in the previous section (*Section 3 Theoretical framework*) and assumptions on the correlations between the trait and the species’ probability of naturalization and invasion (Figure 2). Then, we extended the framework to incorporate variability in the correlations between the trait and the species probabilities to be introduced, naturalize and become invasive, as well as correlations in opposite direction among stages (i.e. trait had a positive correlation with the naturalization probability but negative with the invasion probability). Finally, we exemplified how the framework can be applied to real plant functional traits widely used across invasiveness studies.

**Figure 2.**
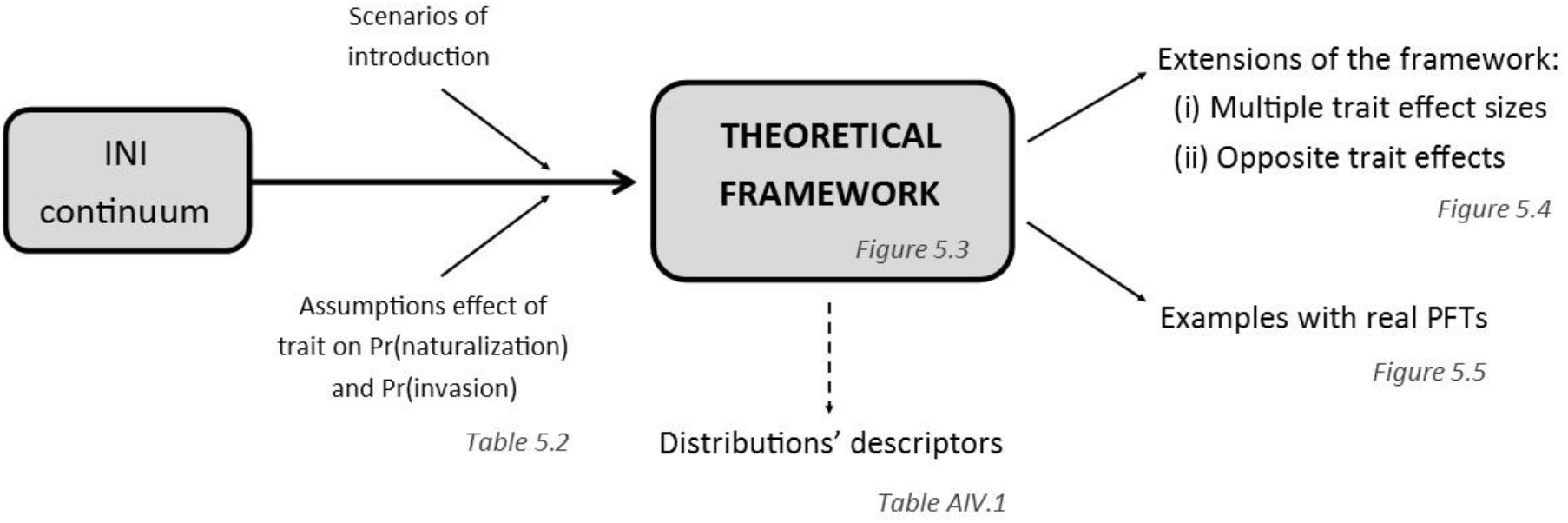
Methodological workflow of Chapter 5. We use a stage-based approach (introduction-naturalization-invasion continuum; Richardson et al. 2000, Blackburn et al. 2011) to develop a theoretical framework of the impact that introduction bias may have on invasion patterns. We propose three different scenarios of introduction -*Random, Biologically biased* and *Human biased*- and make some assumptions on the correlations between a hypothetical trait and the probability of naturalization and invasion. We then extend the examples presented in the hypothetical framework by showing the possible variability within simulations where the correlations between the trait and the probability of naturalization and/or invasion are more or less strong, and when the trait relates to those two stages in opposite directions. Finally, we exemplify how the framework can be applied to real plant functional traits widely used across the invasiveness literature.

### 4.1 Simulation of the theoretical framework

We simulated the distribution of values of a hypothetical continuous trait, ranging from 1 to 100, for the introduction, naturalization and invasion stages. For simplicity, we only present the simulated distribution of values for several possible scenarios for each stage (Figure 3). We assume that the trait either does or does not influence (i.e. correlate with) the probability of introduction, naturalization and/or invasion. When a trait is correlated with naturalization or invasion, effects are always linear, positive, and of similar magnitude.

**Figure 3.**
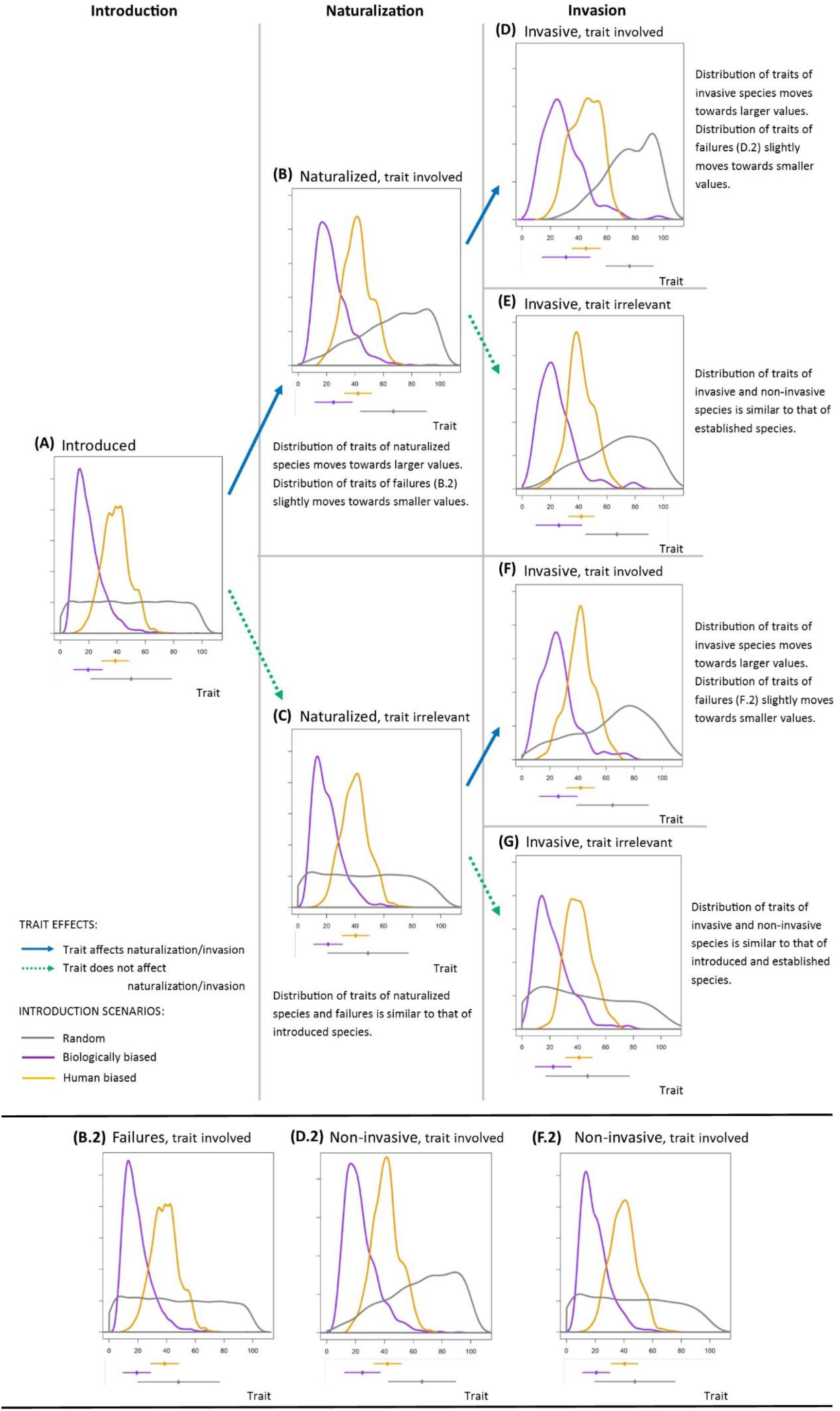
Proposed framework governing the distribution of a hypothetical trait along the introduction-naturalization-invasion continuum for three introduction scenarios: *Random introduction* in grey, *Biologically biased introduction* in purple and *Human biased introduction* in orange. Arrows represent sequential stages and reflect whether the trait is involved in which species successfully reach the next stage (in blue) or not (in green). Trait distribution curves show simulated values for each scenario. Dots and lines below each panel represent the mean and standard deviation of each distribution. Summaries of means, standard deviations and kurtosis values for all possible scenarios are presented in Table A1. If the introduction scenario giving rise to a particular trait distribution of naturalized or invasive species is unknown, and as a result usually assumed to be *Random*, the effect of the trait on species’ success may be overestimated; i.e. the difference between purple or orange lines in panels B to G and grey line in panel A is mistakenly understood as the trait effect. Spurious relationships between traits and species’ success can be minimized by checking whether traits that promotes success - naturalized, invasive species – also correlate in the opposite way with lack of success – failures, non-invasive species (panels B.2, D.2, F.2).

#### 4.1.1 Stage 1: Introduction

We simulated the introduction of 10 000 species along a range of trait values from 1 to 100 (see Panel A2). For the *Random introduction*, we assigned a uniform distribution to the trait values of introduced species. For the *Biologically biased introduction*, we assumed that the trait of the introduced species followed the distribution of the trait for the worldwide pool of species. An inspection of functional traits literature (Table 1) revealed that a lognormal shape was a reasonable assumption for a hypothetical trait, with the distribution being skewed towards small values. Accordingly, we simulated a log-transformed normal distribution with (normal) mean 20 and (normal) standard deviation 10, which ranged from 5.4 (0.5% quantile) to 60.5 (99.5% quantile). For the *Human biased introduction*, we subsampled the *Biologically biased introduction* in a way that the resulting distribution (normal) mean was around 40, instead of 20 (standard deviation was left unchanged), and the range of values went from 16 to 65.9 (0.5% - 99.5% quantiles). This emulates a situation where species with larger (than the mean) values of the hypothetical trait are preferentially introduced from the available worldwide pool; for example, for ornamental purposes, plants with higher above-ground biomass are preferred (Townsley-Brascamp & Marr 1995; Schlaepfer et al. 2010).

**Table 1.**
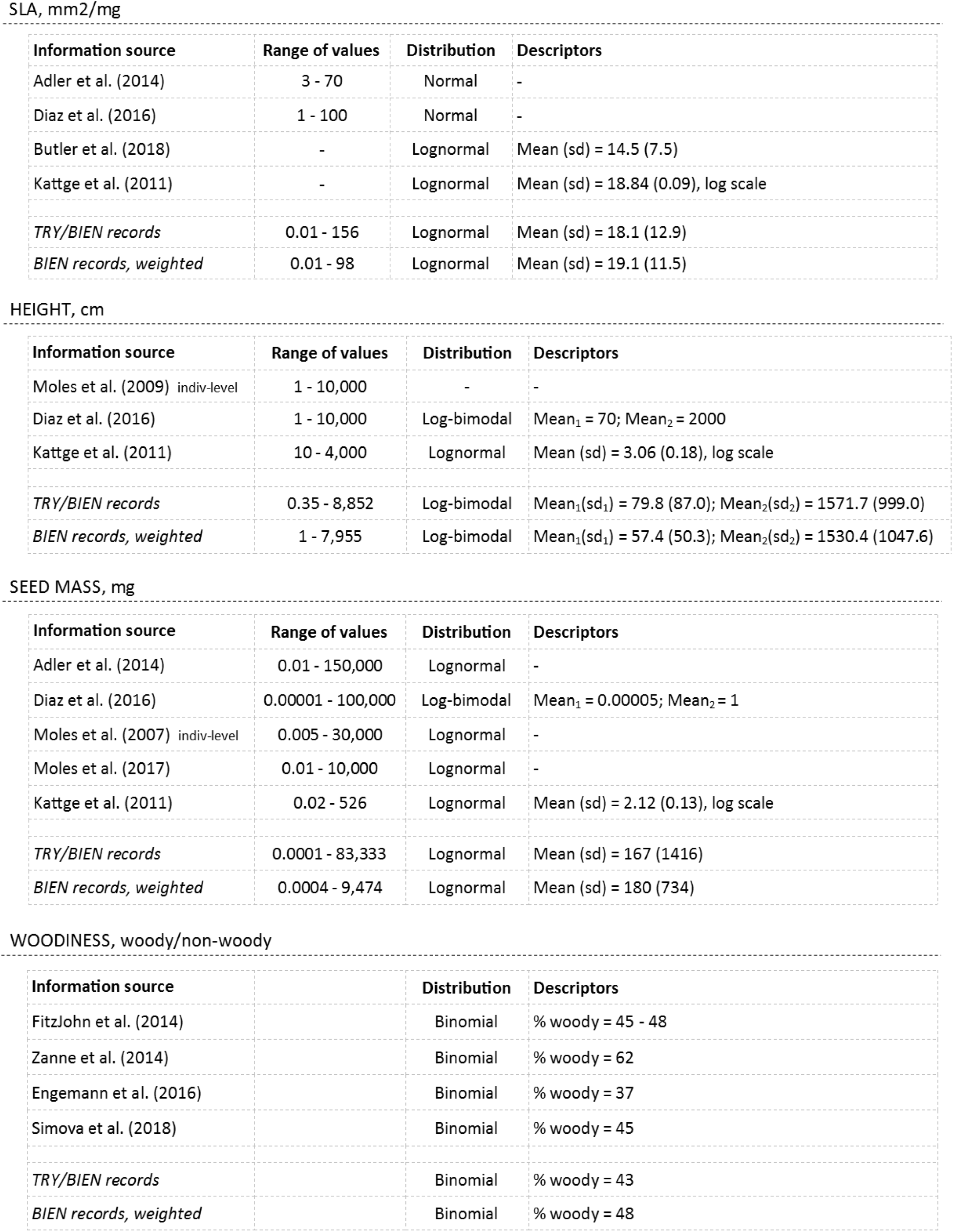
Worldwide biological distribution of plant functional traits’ that are widely-used across the invasiveness literature; SLA, height, seed mass and woodiness. For each trait, their range of values, shape of the distribution and parameters of the distribution (when provided) were extracted from several sources, including peer-reviewed papers and worldwide trait databases. Most papers report trait distributions based on species-level mean values; papers marked with # report trait distributions based on individual-level records. The row ‘TRY/BIEN records’ represents a dataset with the species-level mean trait values available through both the TRY (Kattge et al. 2011) and BIEN (Enquist et al. 2016, Maitner et al. 2018) datasets combined. The row ‘BIEN records, weighted’ represents a dataset with species-level means collated from BIEN database, with species contribution weighted following the species’ worldwide distribution – more widely available species contribute more values to the distribution (see Panel A1).

| Information source | Range of values | Distribution | Descriptors |
| --- | --- | --- | --- |
| Adler et al. (2014) | 3 - 70 | Normal | - |
| Diaz et al. (2016) | 1 - 100 | Normal | - |
| Butler et al. (2018) | - | Lognormal | Mean (sd) = 14.5 (7.5) |
| Kattge et al. (2011) | - | Lognormal | Mean (sd) = 18.84 (0.09), log scale |
| *TRY/BIEN records* | 0.01 - 156 | Lognormal | Mean (sd) = 18.1 (12.9) |
| *BIEN records, weighted* | 0.01 - 98 | Lognormal | Mean (sd) = 19.1 (11.5) |

HEIGHT, cm
| Information source | Range of values | Distribution | Descriptors |
| --- | --- | --- | --- |
| Moles et al. (2009) indiv-level | 1 - 10,000 | - | - |
| Diaz et al. (2016) | 1 - 10,000 | Log-bimodal | Mean<sub>1</sub> = 70; Mean<sub>2</sub> = 2000 |
| Kattge et al. (2011) | 10 - 4,000 | Lognormal | Mean (sd) = 3.06 (0.18), log scale |
| *TRY/BIEN records* | 0.35 - 8,852 | Log-bimodal | Mean<sub>1</sub>(sd<sub>1</sub>) = 79.8 (87.0); Mean<sub>2</sub>(sd<sub>2</sub>) = 1571.7 (999.0) |
| *BIEN records, weighted* | 1 - 7,955 | Log-bimodal | Mean<sub>1</sub>(sd<sub>1</sub>) = 57.4 (50.3); Mean<sub>2</sub>(sd<sub>2</sub>) = 1530.4 (1047.6) |

SEED MASS, mg
| Information source | Range of values | Distribution | Descriptors |
| --- | --- | --- | --- |
| Adler et al. (2014) | 0.01 - 150,000 | Lognormal | - |
| Diaz et al. (2016) | 0.00001 - 100,000 | Log-bimodal | Mean<sub>1</sub> = 0.00005; Mean<sub>2</sub> = 1 |
| Moles et al. (2007) indiv-level | 0.005 - 30,000 | Lognormal | - |
| Moles et al. (2017) | 0.01 - 10,000 | Lognormal | - |
| Kattge et al. (2011) | 0.02 - 526 | Lognormal | Mean (sd) = 2.12 (0.13), log scale |
| *TRY/BIEN records* | 0.0001 - 83,333 | Lognormal | Mean (sd) = 167 (1416) |
| *BIEN records, weighted* | 0.0004 - 9,474 | Lognormal | Mean (sd) = 180 (734) |

WOODINESS, woody/non-woody
| Information source |  | Distribution | Descriptors |
| --- | --- | --- | --- |
| FitzJohn et al. (2014) |  | Binomial | % woody = 45 - 48 |
| Zanne et al. (2014) |  | Binomial | % woody = 62 |
| Engemann et al. (2016) |  | Binomial | % woody = 37 |
| Simova et al. (2018) |  | Binomial | % woody = 45 |
| *TRY/BIEN records* |  | Binomial | % woody = 43 |
| *BIEN records, weighted* |  | Binomial | % woody = 48 |

#### 4.1.2 Stage 2: Naturalization

Second, we fixed the naturalization rate at 10% (1 000 out of the 10 000 introduced species naturalized). Although the tens rule is a doubtful predictor of the percentage of introduced species that will become naturalized (Jeschke et al. 2012), this number agrees with empirical data of plant invasions in Australia (Randall 2012) and New Zealand (Diez et al. 2009). For each introduction scenario, we first assumed that the trait had a linear, positive effect on naturalization, i.e., species with larger trait values had higher probabilities of naturalization. Linear trait effects were calculated without intercepts and were standardized among all values of a given trait, so that the probability to become naturalized increases 100 times from species with trait value 1 to species with trait value 100. Alternatively, for each introduction scenario, we run simulations assuming that naturalization had no relationship with the trait values, and therefore species were randomly drawn from the available pool of 10 000 introduced species (Panel A2). In all cases, species with trait values that were not selected in these simulations (i.e. species from the introduced pool that did not naturalize) were grouped and labelled as Failures.

#### 4.1.3 Stage 3: Invasion

For each naturalization simulation, we fixed the invasion rate at 10% (100 out of 1 000 naturalized species became invasive). Again, species with particular trait values may have an increased probability of invasion or, alternatively, the probability of becoming invasive may be unrelated to the trait. To simulate the former invasive species pool, we assumed that the trait had a linear, positive effect on invasion so that species with larger values had higher probability of becoming invasive. As above, linear trait effects were calculated without intercepts and were standardized among all values of a given trait, so that the probability of invasion increases 100 times from species with trait value 1 to species with trait value 100. The latter pool of invasive species was drawn by randomly subsampling the naturalized pool of species. Species with trait values that were not selected in these simulations (i.e. species from the naturalized pool that did not become invasive) were grouped and labelled as Non-invasive.

### 4.2 Characterization of trait distributions

For each of the 3 proposed introduction scenarios, 13 trait distributions were simulated (1 for the introduced pool, 4 for the naturalized/failures pools, 8 for the invasive/non-invasive pools), resulting in a total of 39 simulations. For each simulated trait distribution, mean, standard deviation and kurtosis were calculated.

### 4.3 Extensions of the framework

After building and examining the base model, we explored how the trait distributions changed if the hypothetical trait had stronger or weaker relationships (i.e. varying effect sizes) with the introduction, naturalization or invasion probabilities. We assumed that all these relationships were positive.

To simulate the consequence of various effect sizes in the *Human biased introduction*, we subsampled the *Biologically biased introduction* three times, fixing the mean of the resulting log-transformed normal distribution around 30, 45 and 60 (normal scale). The resulting three *Human biased introduction* scenarios represent an increasing preference for species with larger and larger trait values out of the available worldwide pool.

Simulations for the naturalization stage assumed trait effect sizes of 0.0025, 0.05 and 0.25 when subsampling each introduction scenario through a loglinear function. Simulations of naturalization drawn from the *Human biased introduction* scenario are based on the medium level of human preference. We used the same approach to run the simulations of the invasion stage.

The second extension explored how opposite effects of the trait on the naturalization and invasion stages affected the trait distributions. We assumed linear positive effects of magnitude 0.005 and loglinear negative effects of magnitude –0.005, which defined opposite patterns of selection across the range of trait values.

### 4.4 Applying the framework to commonly used plant functional traits

We simulated some of the introduction, naturalization and invasion trait distributions for a selection of plant functional traits widely used across the invasion literature: specific leaf area (SLA), height, seed mass and woodiness. In every case, we simulated the introduction of 10 000 species and examined previously published literature to draw the *Random* and *Biologically biased introduction* scenarios as realistically as possible (Table 1). Our simulations assume that the worldwide distributions of SLA and seed mass have a lognormal shape, height has a log-transformed, bimodal distribution and woodiness is a binary trait with woody and non-woody species having similar relative frequencies.

We limited the *Human biased introduction* scenarios to horticultural introductions (Table 2), since this is the most extensively studied pathway across the invasion literature and is considered a major contributor to plant invasions in Australia (Dodd et al. 2015) and worldwide (van Kleunen et al. 2018b). With a few exceptions (Dehnen-Schmutz et al. 2007; van Kleunen et al. 2007), most studies examining horticultural introductions report broad expected relationships between plant characteristics and human preferences for ornamental plants (Table 2, Stage 1: Introduction). As a result, to draw the *Human biased introduction* scenario, we decided on a correlation between the traits and the probability of introduction in agreement with the (limited) available information (Table 2).

**Table 2.**
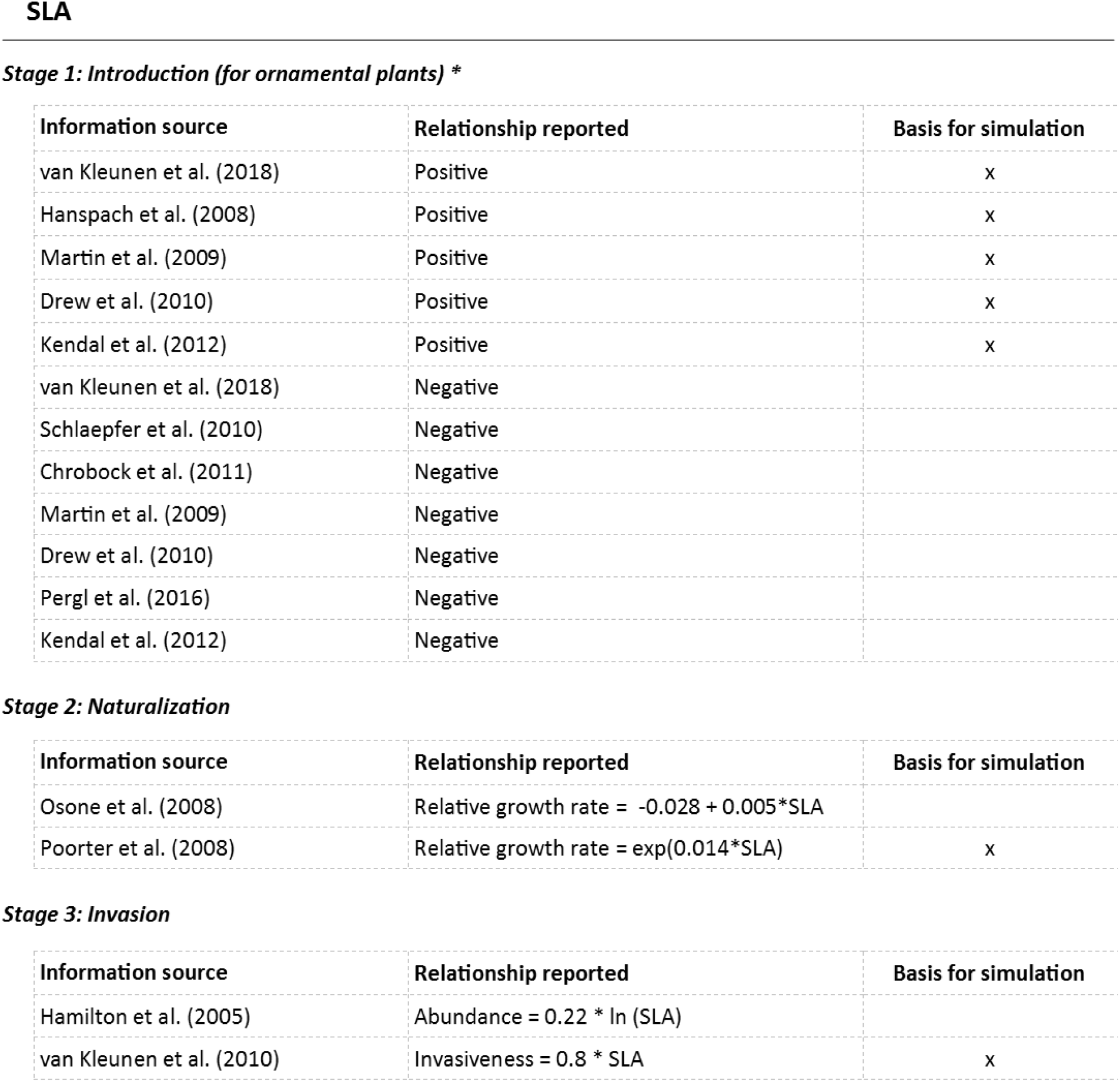
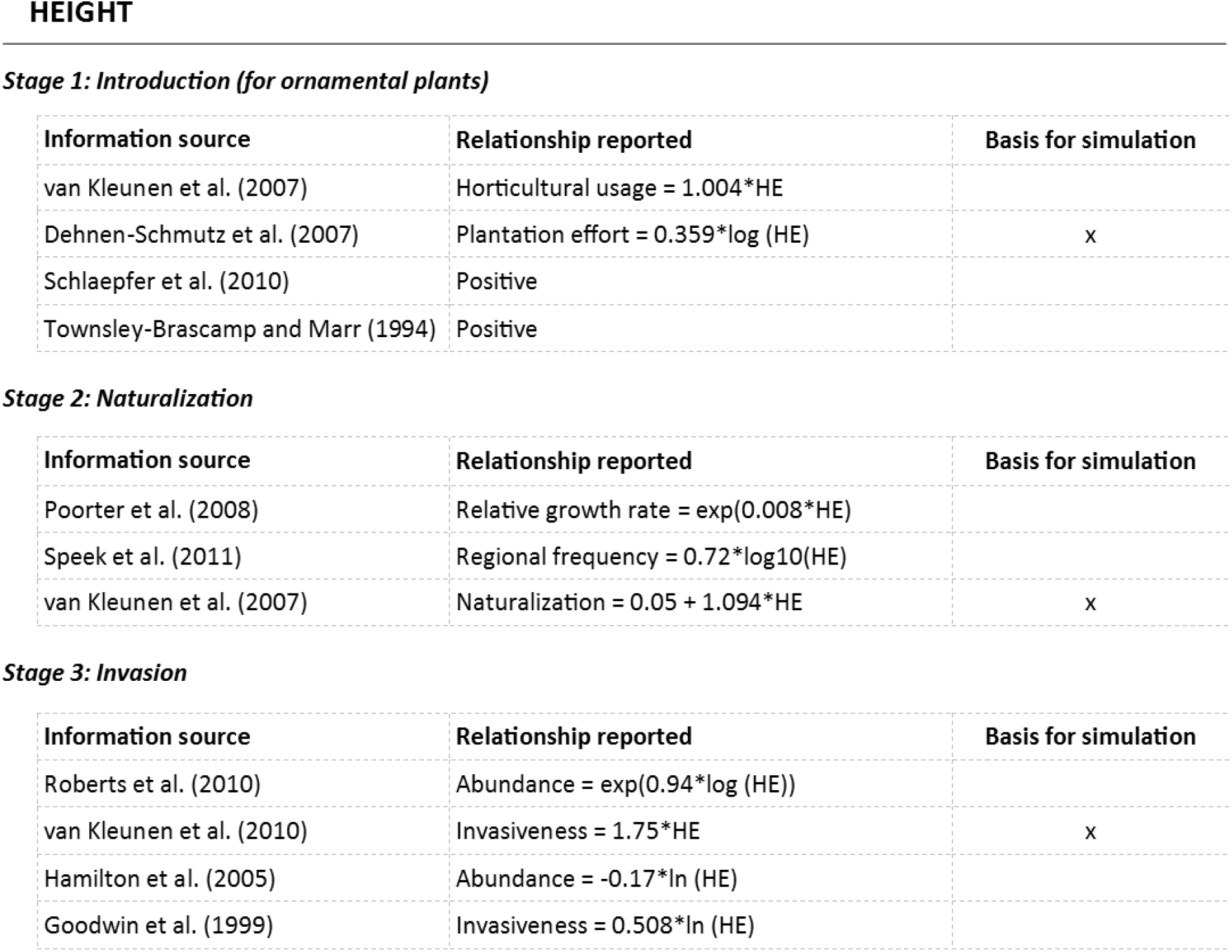
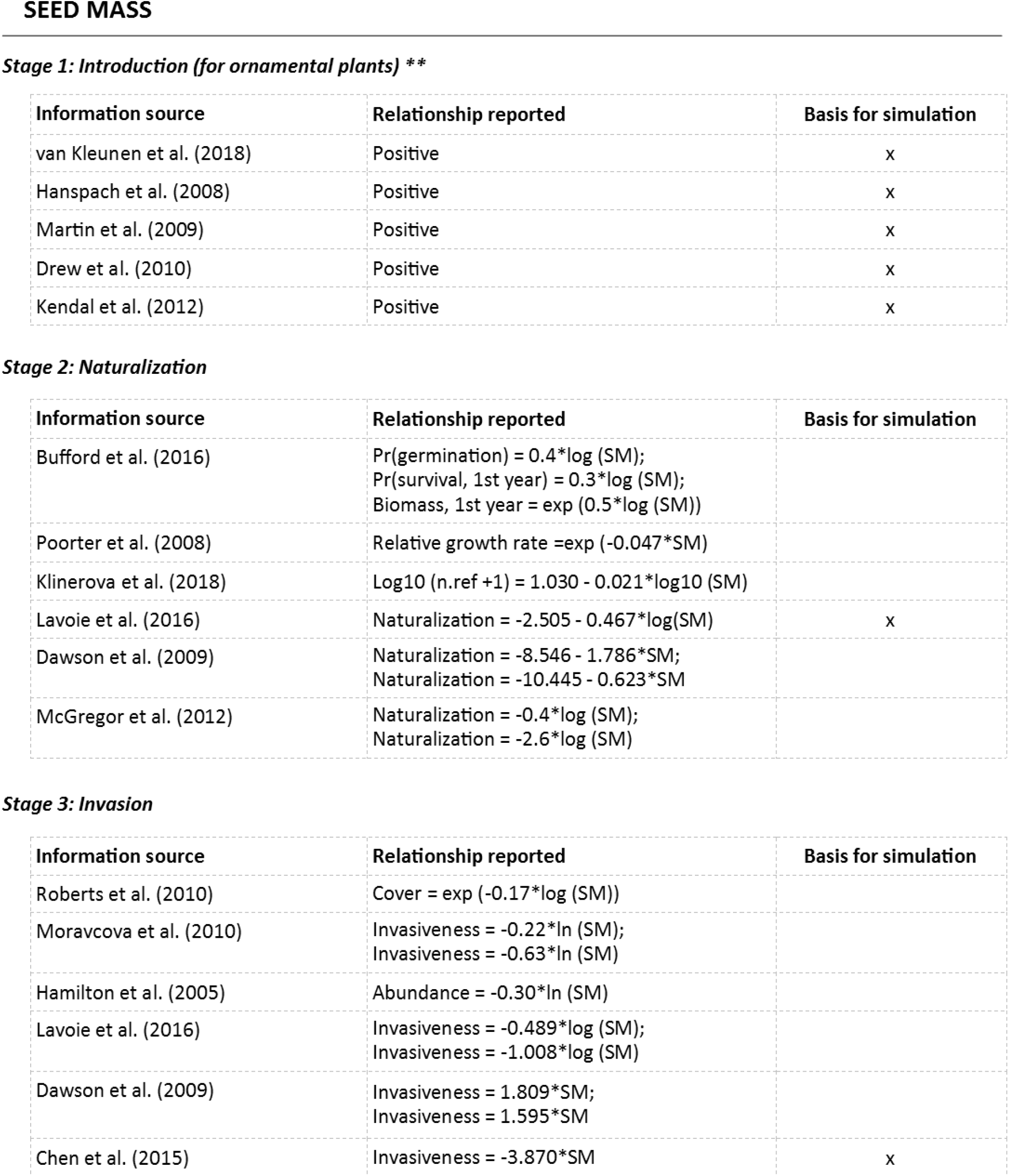
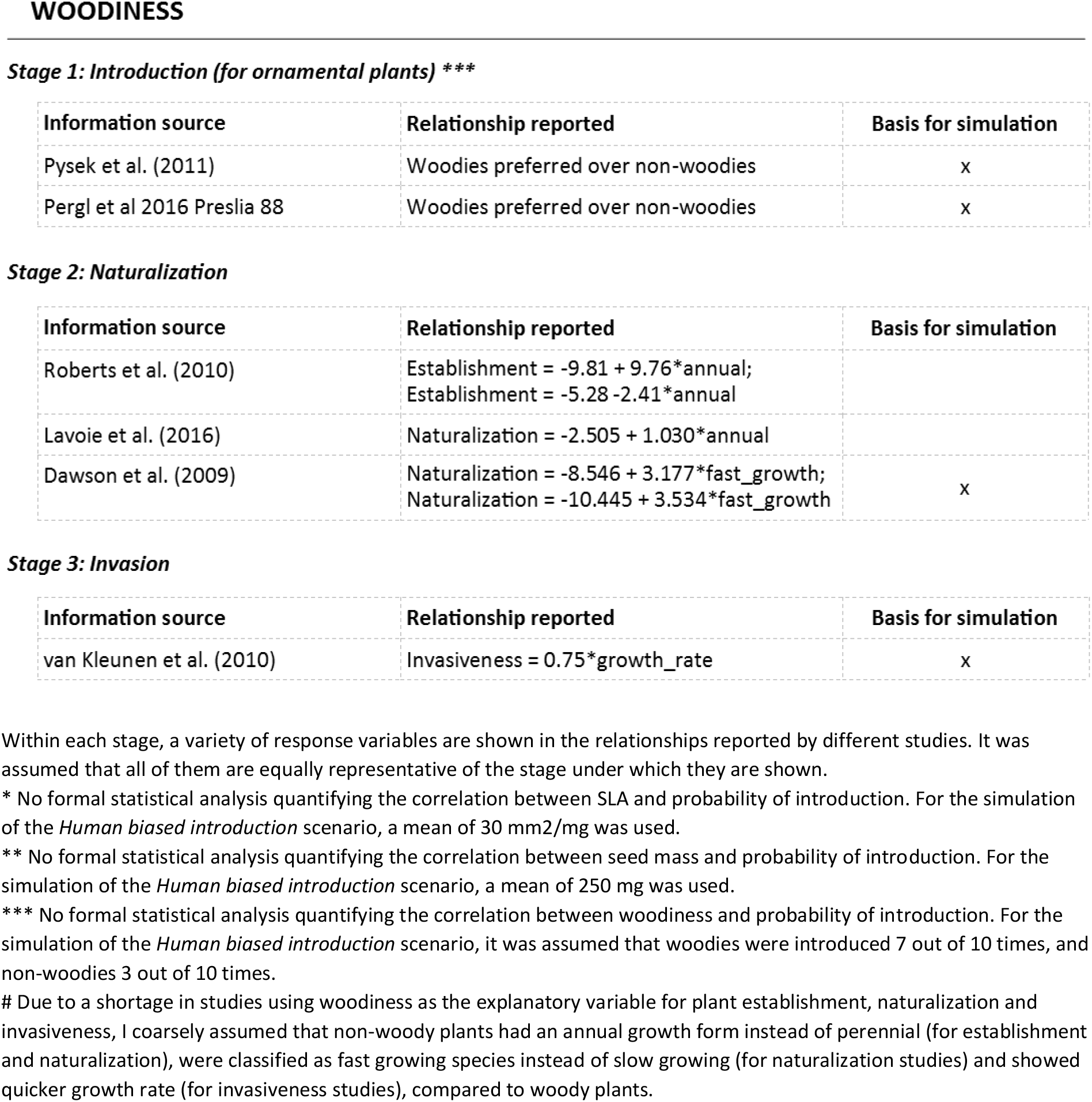
Summary of literature describing the relationship between plant functional traits and human preference of introduction of ornamental plants (Stage 1: Introduction), naturalization success (Stage 2: Naturalization) and invasiveness (Stage 3: Invasion). Trait effect size are included when they are reported in the source; otherwise general trend as reported in the source is presented. Column ‘Basis for simulation’ indicates the source used to define the correlation between the traits and the probabilities of introduction (for the *Human biased introduction* scenario), naturalization and invasion

*Stage 1: Introduction (for ornamental plants) \**
| Information source | Relationship reported | Basis for simulation |
| --- | --- | --- |
| van Kleunen et al. (2018) | Positive | x |
| Hanspach et al. (2008) | Positive | x |
| Martin et al. (2009) | Positive | x |
| Drew et al. (2010) | Positive | x |
| Kendal et al. (2012) | Positive | x |
| van Kleunen et al. (2018) | Negative |  |
| Schlaepfer et al. (2010) | Negative |  |
| Chrobock et al. (2011) | Negative |  |
| Martin et al. (2009) | Negative |  |
| Drew et al. (2010) | Negative |  |
| Pergl et al. (2016) | Negative |  |
| Kendal et al. (2012) | Negative |  |

*Stage 2: Naturalization*
| Information source | Relationship reported | Basis for simulation |
| --- | --- | --- |
| Osone et al. (2008) | Relative growth rate = $-0.028 + 0.005 \cdot \text{SLA}$ | |
| Poorter et al. (2008) | Relative growth rate = $\exp(0.014 \cdot \text{SLA})$ | x |

*Stage 3: Invasion*
| Information source | Relationship reported | Basis for simulation |
| --- | --- | --- |
| Hamilton et al. (2005) | Abundance = $0.22 \cdot \ln(\text{SLA})$ | |
| van Kleunen et al. (2010) | Invasiveness = $0.8 \cdot \text{SLA}$ | x |

HEIGHT *Stage 1: Introduction (for ornamental plants)*
| Information source | Relationship reported | Basis for simulation |
| --- | --- | --- |
| van Kleunen et al. (2007) | Horticultural usage = $1.004 \cdot \text{HE}$ | |
| Dehnen-Schmutz et al. (2007) | Plantation effort = $0.359 \cdot \log(\text{HE})$ | x |
| Schlaepfer et al. (2010) | Positive |  |
| Townsley-Brascamp and Marr (1994) | Positive |  |

*Stage 2: Naturalization*
| Information source | Relationship reported | Basis for simulation |
| --- | --- | --- |
| Poorter et al. (2008) | Relative growth rate = $\exp(0.008 \cdot \text{HE})$ | |
| Speek et al. (2011) | Regional frequency = $0.72 \cdot \log_{10}(\text{HE})$ | |
| van Kleunen et al. (2007) | Naturalization = $0.05 + 1.094 \cdot \text{HE}$ | x |

*Stage 3: Invasion*
| Information source | Relationship reported | Basis for simulation |
| --- | --- | --- |
| Roberts et al. (2010) | Abundance = $\exp(0.94 \cdot \log(\text{HE}))$ | |
| van Kleunen et al. (2010) | Invasiveness = $1.75 \cdot \text{HE}$ | x |
| Hamilton et al. (2005) | Abundance = $-0.17 \cdot \ln(\text{HE})$ | |
| Goodwin et al. (1999) | Invasiveness = $0.508 \cdot \ln(\text{HE})$ | |

SEED MASS Stage 1: Introduction (for ornamental plants) \*\*
| Information source | Relationship reported | Basis for simulation |
| --- | --- | --- |
| van Kleunen et al. (2018) | Positive | x |
| Hanspach et al. (2008) | Positive | x |
| Martin et al. (2009) | Positive | x |
| Drew et al. (2010) | Positive | x |
| Kendal et al. (2012) | Positive | x |

Stage 2: Naturalization
| Information source | Relationship reported | Basis for simulation |
| --- | --- | --- |
| Bufford et al. (2016) | Pr(germination) = $0.4 \cdot \log(\text{SM})$;<br>Pr(survival, 1st year) = $0.3 \cdot \log(\text{SM})$;<br>Biomass, 1st year = $\exp(0.5 \cdot \log(\text{SM}))$ | |
| Poorter et al. (2008) | Relative growth rate = $\exp(-0.047 \cdot \text{SM})$ | |
| Klinerova et al. (2018) | $\log_{10}(\text{n.ref} + 1) = 1.030 - 0.021 \cdot \log_{10}(\text{SM})$ | |
| Lavoie et al. (2016) | Naturalization = $-2.505 - 0.467 \cdot \log(\text{SM})$ | x |
| Dawson et al. (2009) | Naturalization = $-8.546 - 1.786 \cdot \text{SM}$;<br>Naturalization = $-10.445 - 0.623 \cdot \text{SM}$ | |
| McGregor et al. (2012) | Naturalization = $-0.4 \cdot \log(\text{SM})$;<br>Naturalization = $-2.6 \cdot \log(\text{SM})$ | |

Stage 3: Invasion
| Information source | Relationship reported | Basis for simulation |
| --- | --- | --- |
| Roberts et al. (2010) | Cover = $\exp(-0.17 \cdot \log(\text{SM}))$ | |
| Moravcova et al. (2010) | Invasiveness = $-0.22 \cdot \ln(\text{SM})$;<br>Invasiveness = $-0.63 \cdot \ln(\text{SM})$ | |
| Hamilton et al. (2005) | Abundance = $-0.30 \cdot \ln(\text{SM})$ | |
| Lavoie et al. (2016) | Invasiveness = $-0.489 \cdot \log(\text{SM})$;<br>Invasiveness = $-1.008 \cdot \log(\text{SM})$ | |
| Dawson et al. (2009) | Invasiveness = $1.809 \cdot \text{SM}$;<br>Invasiveness = $1.595 \cdot \text{SM}$ | |
| Chen et al. (2015) | Invasiveness = $-3.870 \cdot \text{SM}$ | x |

WOODINESS Stage 1: Introduction (for ornamental plants) \*\*\*
| Information source | Relationship reported | Basis for simulation |
| --- | --- | --- |
| Pysek et al. (2011) | Woodies preferred over non-woodies | x |
| Pergl et al 2016 Preslia 88 | Woodies preferred over non-woodies | x |

Stage 2: Naturalization
| Information source | Relationship reported | Basis for simulation |
| --- | --- | --- |
| Roberts et al. (2010) | Establishment = $-9.81 + 9.76 \cdot \text{annual}$;<br>Establishment = $-5.28 - 2.41 \cdot \text{annual}$ | |
| Lavoie et al. (2016) | Naturalization = $-2.505 + 1.030 \cdot \text{annual}$ | |
| Dawson et al. (2009) | Naturalization = $-8.546 + 3.177 \cdot \text{fast\_growth}$;<br>Naturalization = $-10.445 + 3.534 \cdot \text{fast\_growth}$ | x |

Stage 3: Invasion
| Information source | Relationship reported | Basis for simulation |
| --- | --- | --- |
| van Kleunen et al. (2010) | Invasiveness = $0.75 \cdot \text{growth\_rate}$ | x |

Within each stage, a variety of response variables are shown in the relationships reported by different studies. It was assumed that all of them are equally representative of the stage under which they are shown.
\* No formal statistical analysis quantifying the correlation between SLA and probability of introduction. For the simulation of the *Human biased introduction* scenario, a mean of 30 mm<sup>2</sup>/mg was used.
\*\* No formal statistical analysis quantifying the correlation between seed mass and probability of introduction. For the simulation of the *Human biased introduction* scenario, a mean of 250 mg was used.
\*\*\* No formal statistical analysis quantifying the correlation between woodiness and probability of introduction. For the simulation of the *Human biased introduction* scenario, it was assumed that woodies were introduced 7 out of 10 times, and non-woodies 3 out of 10 times.

The relationships between traits and naturalization probability were extracted from seed addition and other field experiments recording plant establishment (Table 2, Stage 2: Naturalization). When possible, we avoided relying on invasion literature for the naturalization stage because introduction bias is often overlooked when examining traits of non-native species. We did, however, rely on invasion literature to determine the effect of the traits on the subsequent stage—the probability of naturalized species becoming invasive (Table 2, Stage 3: Invasion). Consistent with our hypothetical framework, we assumed that only 10% of the species pool was able to reach the subsequent stage of the introduction-naturalization-invasion continuum.

## 5 Results

The assumptions made for the introduction scenarios have an impact on the shape of the simulated trait distributions of both naturalized and invasive species. Evaluating the trait patterns observed on naturalized and invasive species while overlooking the possibility that introduction may be biased towards particular values (Figure 3 A - purple and orange). Therefore, the assumption it happens randomly (Figure 3 A - grey), may lead to an overestimation of the implication of the trait on naturalization and invasion success. The overestimation appears as the result of overlooking that the difference between the trait distributions of naturalized and invasive species (Figure 3 – purple and orange) and the *Random introduction* scenario (Figure 3 A – grey) is the combination of any existing trait effect on naturalization and/or invasion and any existing bias in the species introduction.

### 5.1 The theoretical framework

The distribution of trait values along the introduction-naturalization-invasion continuum only changes when the trait has an effect on the probabilities of species becoming naturalized and/or invasive (Figure 3 B, D, F), as reflected by the different mean, standard deviation and kurtosis of the trait distributions simulated for introduced, naturalized and invasive species for each introduction scenario (Table A1 A, B, D, F). Regardless of the introduction scenario (Figure 3 A), the differences between introduced and invasive species increase when trait effects accumulate across stages; e.g. the trait increases the naturalization and invasion probability (Figure 3 D) compared to scenarios where the trait affects either naturalization (Figure 3 E) or invasion (Figure 3 F) but not both. However, the differences in trait distributions among stages (for a particular trait effect size) are smaller as the bias in the introduction becomes larger, meaning that trait differences among introduced, naturalized and invasive species are larger for species introduced randomly and smaller for species introduced through the assumed hypothetical human pathway. In general, naturalized and invasive species drawn from the randomly introduced species pool (Figure 3 – grey) have fairly flat distributions, with high means and large standard deviations (Table A1, *Random introduction* column). By contrast, those drawn from a *biologically biased introduced* species pool (Figure 3 - purple) have the most-peaked distributions with the smallest mean values (Table A1, *Biologically biased introduction* column), and those drawn from a pool of introduced species through the assumed, hypothetical human introduction pathway (Figure 3 - orange) sit between the other two scenarios and have a normal-shaped distribution that changes little across stages (Table A1, *Human biased introduction* column). No differences in the trait distribution are found between consecutive stages when the trait has no effect on the naturalization or invasion probability (Figure 3 C, E, G).

Within a single stage of the framework, successful (i.e. naturalized, invasive) and unsuccessful (i.e. failures, non-invasive) species will show different trait distributions only if the trait is involved in naturalization (Figure 3 B vs B.2) or invasion (Figure 3 D vs. D.2, F vs. F.2). If the trait is not involved in species’ success, both successful and unsuccessful species will mimic the trait distribution of species in the previous stage (Figure 3 C, E, G). Even if the trait is involved in species’ success, unsuccessful species (Figure 3 B.2, D.2, F.2) still resemble the distribution of the species at the previous stage because the simulations assumed that most introduced/naturalized species (90%) fail to become naturalized/invasive (*Section 4*.*1 Simulation of the theoretical framework*).

### 5.2 Extensions of the framework

Regardless of the magnitude of trait effects on introduction, naturalization and invasion assumed in our framework (i.e. strength of the correlation between the trait and the probability of being introduced, naturalize or become invasive), the relative differences among the distributions of subsequent stages remained very similar, mostly mimicking the differences among introduction scenarios (Figure 4i). The exception was the distribution of naturalized and invasive species after *Random introduction*, which moved towards extremely high trait values when the effect size was high (e.g. Figure 4i D).

**Figure 4.**
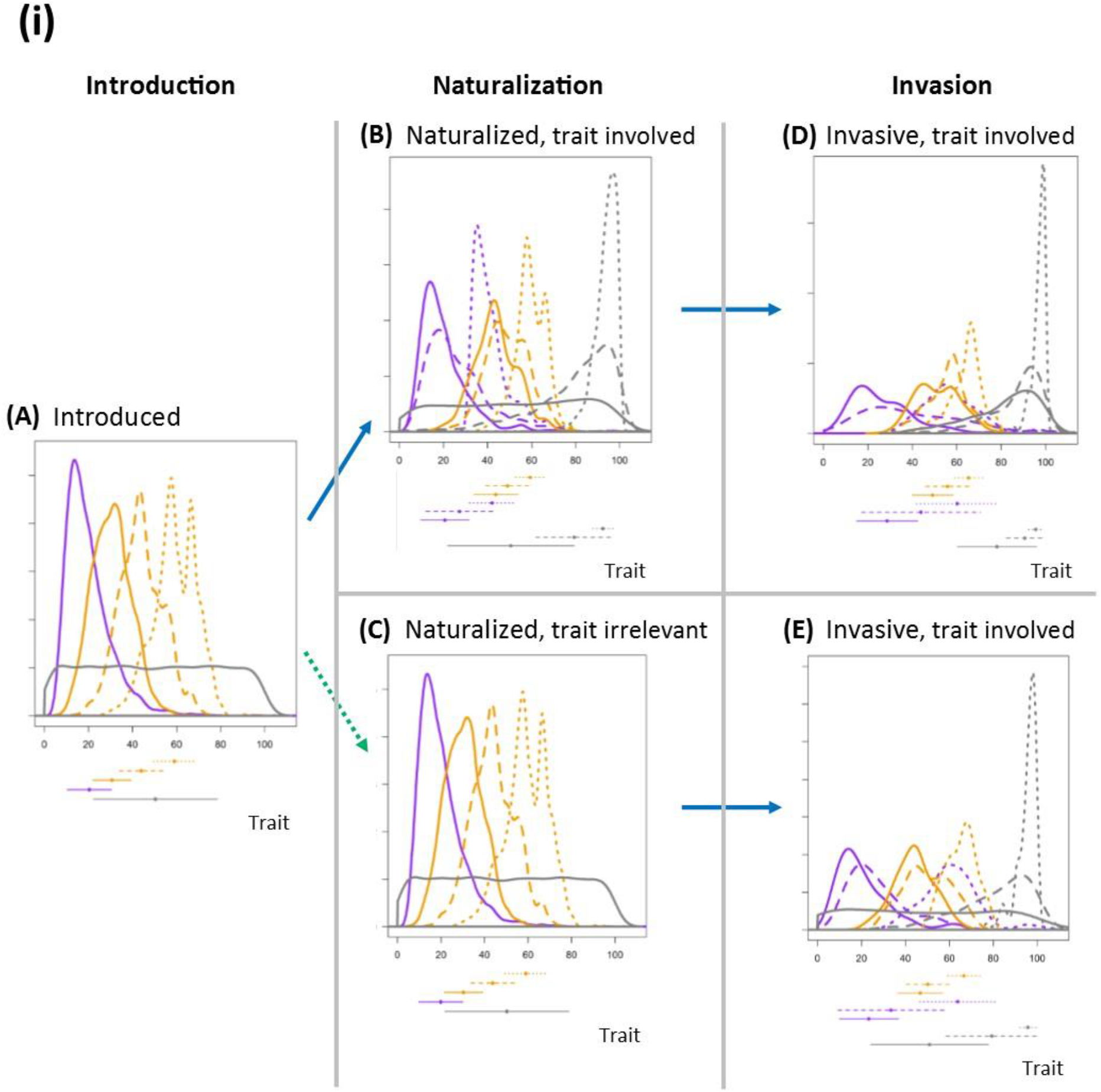
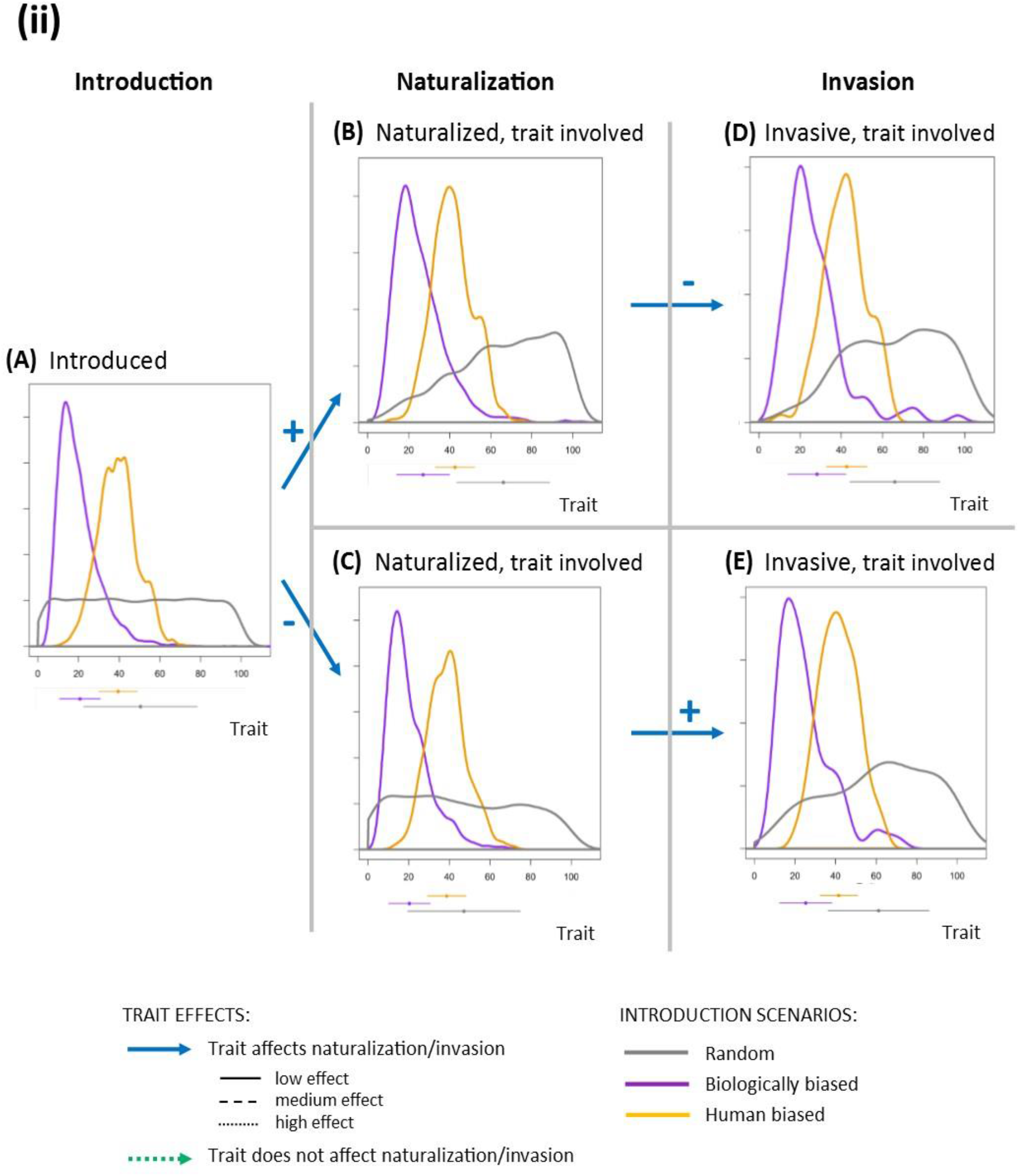
**(i)** Extension of some of the scenarios presented in the proposed framework including low, medium and high trait effect size on introduction, naturalization and invasion. The effects are assumed to be positive. Note that higher effects shift distributions towards larger trait values. Within a single stage, distributions resulting from different combinations of introduction bias and trait effect size show large overlaps. **(ii)** Extension of some scenarios presented in the proposed framework including a combination of positive and negative effects of the trait on naturalization and invasion. For each introduction scenario, distributions of traits of invasive species are similar regardless of the order of positive and negative trait effects on naturalization and invasion.

Similar trait distributions for successful species at a given stage may arise from the combination of different assumptions regarding the degree of introduction bias and the strength of the correlation between the trait and the species’ success probability. For example, invasive species drawn from *Biologically biased* and *Human biased introduction* scenarios showed similar trait distributions when the trait strongly and weakly affected the probability of success, respectively (Figure 4i D, dotted purple vs. solid orange). For a given introduction scenario, invasive species also showed similar trait distributions when opposite trait effects on naturalization and invasion probabilities were assumed (Figure 4ii D-E), even though naturalized species did not (Figure 4ii B-C).

### 5.3 Applying the framework to commonly used plant functional traits

Distributions of SLA, height, seed mass and woodiness for naturalized and invasive plants varied among introduction scenarios (Figure 5). In general, for continuous traits, the overlap among simulated distributions for a single stage was low (Figure 5, note log-scaled axes). Simulations based on the *Random introduction* scenario contained many trait values that, based on the traits’ worldwide biological distributions reported in the literature (Table 1), have low availability. For example, SLA has a normal distribution, and therefore few species will show very high values of this trait worldwide; however, a *Random introduction* assumes that species with low, medium and high SLA values are introduced in a similar fashion. As a result, naturalized and invasive species’ trait distributions are highly skewed towards species with high values of the continuous traits, compared to simulations based on the *Biologically biased and Human biased introduction* scenarios. The assumption that introduction was biased towards species with high values - within the biological availability of species - due to human activities (Table 2) resulted in trait distributions of naturalized and invasive species to have a narrower range, limited to the upper range of trait values, in comparison with those drawn from the *Biologically biased introduction*. In the case of woodiness, trait distributions based on the *Random* and *Biologically biased introduction* scenarios looked similar - an assumption of a 50:50 ratio of woody and non-woody plants for the *Biologically biased introduction* seems reasonable (Table 2).

**Figure 5.**
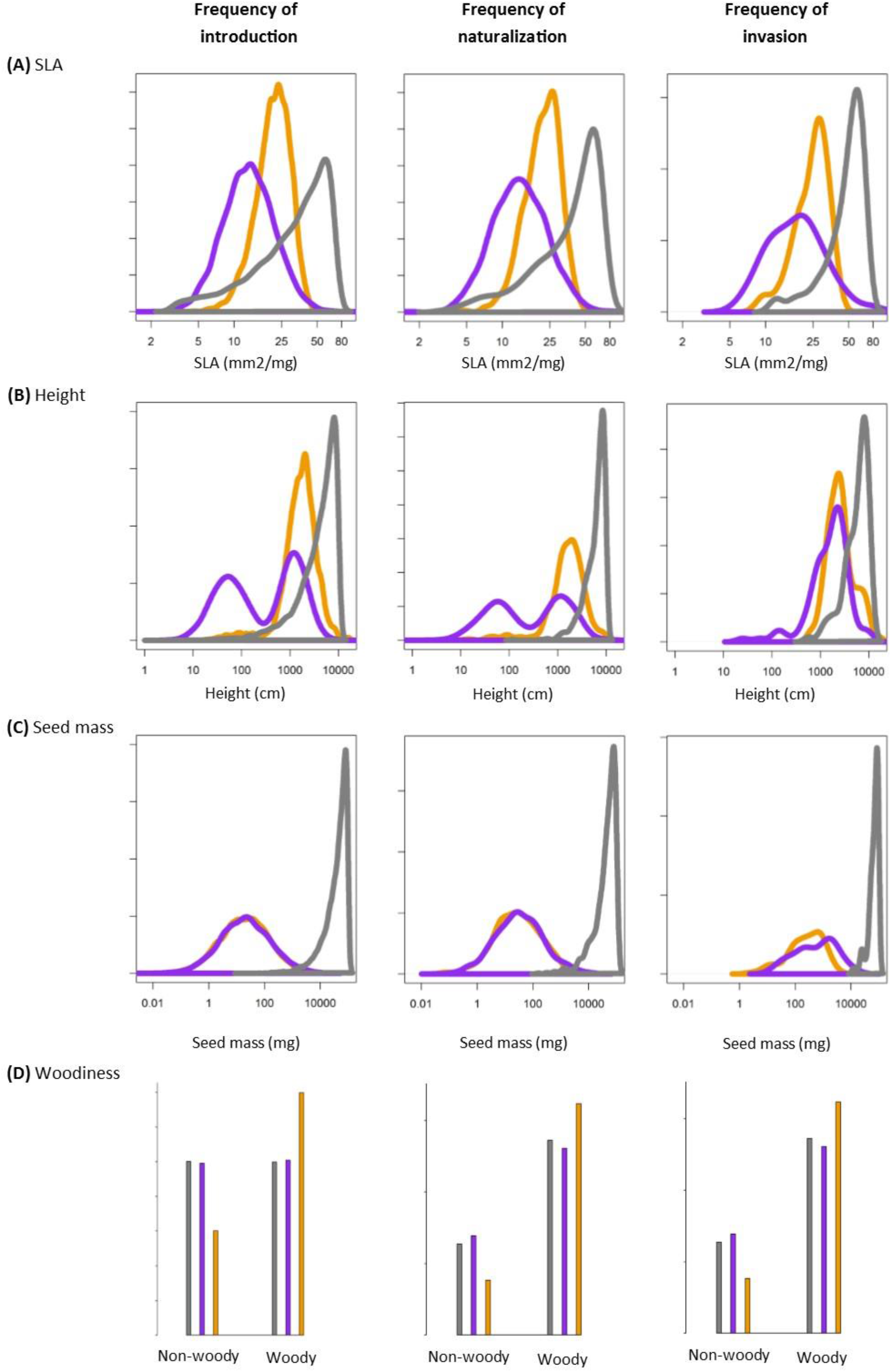
Hypothetical distribution of specific leaf area (SLA), height, seed mass and woodiness for introduced, naturalized and invasive species, assuming *Random* (in grey), *Biologically biased* (purple) and *Human biased* (orange) *introduction* scenarios. The assumed effect of traits on the probability of introduction, (for the *Human biased introduction* scenario), naturalization and invasion follow published literature (Table 2). The group of species, and therefore the distribution of trait values, that are introduced in the first place highly influences the group of species that ultimately become invasive. The differences among distributions simulated for invasive species under the three introduction scenarios suggest that invasiveness studies overlooking any introduction bias that these traits may show may lead to erroneous conclusions regarding the traits’ correlations with plant invasiveness.

## 6 Discussion

### 6.1 Benefits of the theoretical framework

Through use of a theoretical framework and simulation models, in this paper we have demonstrated how different functional biases in the pool of introduced species can affect the trait distributions of successful species in subsequent stages along the introduction-naturalization-invasion continuum (i.e. naturalization and invasion). Overlooking introduction bias can potentially lead to erroneous conclusions about traits associated with naturalization and invasion (Blackburn & Jeschke 2009; Kueffer et al. 2013), especially in studies that investigate functional patterns of naturalized species but ignore the introduced species that have failed to naturalize, and in studies that investigate traits correlated with invasive species by comparing them with native species. Our theoretical framework is a starting point to explore likely ways that trait-based invasion studies overlooking the introduction stage may report results that conflate trait and introduction bias effects.

Functional studies of invasion rarely account for the whole introduced pool of species. They usually implicitly assume that species introduction is independent from the trait under examination and that all values of said trait have similar probability of being introduced (Figure 3 A - grey). As a result, any departure from a random distribution found in the trait of naturalized or invasive species (Figure 3 B to G – purple and orange) may be misinterpreted as the effect of the trait alone on naturalization or invasion, respectively, regardless of the true magnitude of the trait effect. The extent to which a trait effect can be under or overestimated will depend on the true magnitude of the effect and the type of introduction bias. The overestimation of trait effects on naturalization and invasion due to the assumption that introduction happens randomly will increase for those cases where the introduced species show a large functional bias (Figure 3 A Human bias introduction).

Inspecting the functional identity of unsuccessful species can reveal any functional bias in the introduction and clarify whether the trait distributions of naturalized and invasive species respond to a true ecological benefit of having particular values of the trait under examination (e.g. Figure 3 B vs. B.2) or they are an artefact of introduction (e.g. Figure 3 C). A true correlation between a trait and species’ ability to reach a given stage can be confirmed by finding the opposite effect on species’ that have failed to reach that stage; that is the traits must have opposite effects on established species vs. failures (for the naturalization stage) or on invasive vs. non-invasive species (for the invasion stage). While studies focused on the invasion stage commonly include information on the whole pool of naturalized species, by comparing successful (i.e. invasive) and not successful (i.e. non-invasive) species (Chen et al. 2015; Gallagher et al. 2015; Lavoie et al. 2016), knowledge on the whole pool of introduced species in an area is usually not available (e.g. Hamilton et al. 2005). As a result, studies focused on the naturalization stage are often limited to the group of successful (i.e. naturalized) species (but see Lavoie et al. 2016; Dawson et al. 2009; McGregor et al. 2012), which may lead to spurious results on the role of traits. Likewise, studies that aim to determine traits of invasive species by comparing them to native species, yet overlooking non-invasive species (Funk & Throop 2010; Godoy et al. 2012; but see Bezeng et al. 2015), may also conflate effects of traits and introduction bias. The difference between invasive and native trait distributions will increase as the introduced species are biased toward extreme trait values or if they present values novel to the community, which may increase their ability to fill empty niches. Our framework suggests that attempts to identify traits that facilitate invasion need to test the traits’ relationship with both invasive and non-invasive species.

The extreme trait distributions of naturalized and invasive species found under the *Random introduction* scenario when the theoretical framework was applied to real plant functional traits (Figure 5) suggests that random introductions are unlikely. If traits were randomly introduced, effect sizes regularly reported in the literature (and used to build our simulations; Table 2) would lead to invasive species showing a narrow range of quite extreme values of their traits (Figure 5 - grey), which is not usually reported in the literature.

### 6.2 Biological bias, preferential introduction and trait-based studies

*Biologically biased introductions* are expected to be characterised by the same taxonomical and functional biases found in the worldwide biota. Plant families that include more species, as well as species that have larger geographic ranges, are expected to be introduced more often. For example, large plant families, such as *Poaceae, Asteraceae* or *Fabaceae*, are often identified as important contributors of invasive species worldwide (see Diez et al. 2009; Dodd et al. 2015 for examples in Australia). Following their worldwide availability (Moles et al, 2007), herbs are expected to be introduced more often, and climbers less often, than other growth forms. However, woody and non-woody species should be introduced in a similar fashion (FitzJohn et al. 2014). Assuming that species from different regions have similar probability to be transported, biological biased introductions worldwide are expected to show similar biases. Our theoretical framework shows a biological bias towards small values of the trait.

*Human biased introductions*, on the other hand, are expected to highly depend on the dominant human activities in the area under examination. Our framework shows scenarios of naturalization and invasion where traits with positive effects on species’ success are preferentially introduced (Figure 3 B, D, F - orange). In general, it is expected that the preselection and movement of species with functional traits valuable for human activity exacerbates the risk of invasion. For example, large quantities of introduced colonizers around urban environments, which are highly disturbed, can potentially lead to higher invasion levels than introduction of random plant types (Martin et al. 2009). Similarly, the introduction of pasture plants with characteristics similar to those of good invaders increases the risk of escape and establishment of exotic species outside agricultural areas (Lonsdale 1994; Driscoll et al. 2014).

Most plant introductions occur either as horticultural escapees or through release of commodities, including deliberate (e.g. plants used to prevent soil erosion or seed trade) and accidental (e.g. agricultural weeds) introductions (Kowarik & von der Lippe 2007; Hulme et al. 2008). So far, most work on functional bias and its consequences for invasion patterns has focused on the horticultural introduction pathway (Kowarik & von der Lippe 2007; Pergl et al. 2017). The types of ornamental plants that are more highly valued, and therefore preferentially introduced, have quick life cycles, generalist requirements, are resistant to harsh environmental conditions and require little maintenance (Hanspach et al. 2008; Martin et al. 2009; Drew et al. 2010; Schlaepfer et al. 2010; van Kleunen et al. 2018b). Even when introductions are accidental, some form of functional bias is expected among the introduced pool of species (Pyšek et al. 2011). One of the first challenges to understand how introduction bias influences invasiveness studies is to translate those generalized patterns of human preferences into particular trait biases. For example, preference for resistant plants under harsh conditions and those with low maintenance requirements may suggest a bias towards species with large seed mass (Leishman et al. 2000; Muller-Landau 2010) and low SLA (Westoby et al. 2002). By contrast, if plants are not introduced as seeds but as established tubestock, plants with quick growth - i.e. small seeds and high SLA - may be preferred (Reich 2014; Moles 2018). Both the traits displaying biases and the nature of these biases depend on the particular introduction pathway under examination (Wilson et al. 2009b).

### 6.3 The way forward

Our hypothetical framework has shown that disregarding human-driven preselection of particular types of introduced species may lead to erroneous conclusions about the causes and correlates of invasion success in plants. Traits that have been introduced more often can be mistakenly interpreted as most likely to adapt to local conditions (as pointed out by Hanspach et al. 2008) or associated with rapid evolution in novel environments (as pointed out by Kitajima et al. 2006 and Chrobock et al. 2011). Future studies of invasion would benefit from critical thinking about how their results may be influenced by a stronger presence of particular forms of successful invaders due to biased introductions (Knapp & Kühn 2012; Buckley & Catford 2016). The challenge of doing so is not only a lack of information on failures, but also differences in the direction and magnitude of trait effects at different stages of the continuum, which depend on the trait under examination and the context of a given study (Dawson et al. 2009). Trade-offs among positive, negative and neutral trait-stage relationships can lead to fading effects of introduction bias or near-zero net effects of traits on invasion (Figure 4ii).

The most suitable approach to validate our framework, and hence how functional biases in the introduced pool may cascade down to successful species in the following stages, is for trait-based studies to evaluate the degree of success of species at multiple stages of invasion (including introduction). Despite their potential value to understand invasion drivers, rarely are records of failed introductions kept. If information on failures has been recorded, the effect of traits on naturalization can be confirmed by checking whether they are related, in an opposite way, with species that fail to naturalize (Figure 3 B vs. B.2) (Diez et al. 2009; Zenni & Nuñez 2013). If records of failures are not available, it may still be possible to indirectly assess the influence of introduction bias.

The correlation between traits and naturalization can be studied independently for species introduced through different pathways; all else being equal, different trait effects across multiple pathways will give an approximate idea of how introductions are biased relative to one another (Kueffer et al. 2013). Introduction bias may be indirectly accounted for in trait-based invasiveness studies by including the interactive effect of species’ traits and species’ introduction frequencies (i.e. propagule pressure), in addition to the additive effects of both variables (Maurel et al. 2016). Although such an approach would not directly speak to constraints on trait values in the introduced pool of species, it should distinguish cases where the trait has an effect on invasiveness from cases where the trait effect is an artefact of high introduction rates of species with those trait values. If none of these approaches are viable because information on neither introduction pathways nor propagule pressure exists, an informal examination of the worldwide or regional distribution of the traits under consideration may provide some perspective regarding the relative abundance (and therefore, availability) of trait values with the potential to be introduced.

## 7 Acknowledgements

The authors thank the Australian Wildlife Society, the Centre of Excellence in Environmental Decisions (CEED), the School of BioSciences, the Botany Foundation and the Australian Research Council for funding. Rod Randall, Mark van Kleunen and Angela Moles kindly provided comments on the manuscript. The dataset associated to Randall (2012) *A Global Compendium of Weeds* was kindly provided by Rod Randall and used to inform some of the simulations.

## References

Adler P. B., Salguero-Gómez R., Compagnoni A., Hsu J. S., Ray-Mukherjee J., Mbeau-Ache C. & Franco M. (2014) Functional traits explain variation in plant life history strategies. Proceedings of the National Academy of Sciences 111, 740–5.

Bezeng S. B., Davies J. T., Yessoufou K., Maurin O. & Van der Bank M. (2015) Revisiting Darwin’s naturalization conundrum: explaining invasion success of non-native trees and shrubs in southern Africa. Journal of Ecology 103, 871–9.

Blackburn T. M., Dyer E., Su S. & Cassey P. (2015) Long after the event, or four things we (should) know about bird invasions. Journal of Ornithology 156, 15–25.

Blackburn T.M. & Jeschke J. M. (2009) Invasion success and threat status: two sides of a different coin? Ecography 32, 83–8.

Blackburn T. M., Pyšek P., Bacher S., Carlton J. T., Duncan R. P., Jarošík V., Wilson J.R.U. & Richardson D. M. (2011) A proposed unified framework for biological invasions. Trends in Ecology & Evolution 26, 333–9.

Buckley Y.M. & Catford J. (2016) Does the biogeographic origin of species matter? Ecological effects of native and non-native species and the use of origin to guide management. Journal of Ecology 104, 4–17.

Bufford J. L., Lurie M.H. & Daehler C. C. (2016) Biotic resistance to tropical ornamental invasion. Journal of Ecology 104, 518–30.

Butler E. E., Datta A., Flores-Moreno H., Chen M., Wythers K. R., Fazayeli F.…, & Reich P. B. (2018) Mapping local and global variability in plant trait distributions. Proceedings of the National Academy of Sciences 114, E10937–E46.

Catford J. A., Baumgartner J. B., Vesk P. A., White M. D., Buckley Y.M. & McCarthy M. A. (2016) Disentangling the four demographic dimensions of species invasiveness. Journal of Ecology 104, 1745–58.

Chen L., Peng S. & Yang B. (2015) Predicting alien herb invasion with machine learning models: biogeographical and life-history traits both matter. Biological Invasions 17, 2187–98.

Chrobock T., Kempel A., Fischer Fischer. & van Kleunen M. (2011) Introduction bias: Cultivated alien plant species germinate faster and more abundantly than native species in Switzerland. Basic and Applied Ecology 12, 244–50.

Colautti R. I., Grigorovich I.A. & MacIsaac H. J. (2006) Propagule pressure: a null model for biological invasions. Biological Invasions 8, 1023–37.

Dawson W., Burslem D.F.R.P. & Hulme P. E. (2009) Factors explaining alien plant invasion success in a tropical ecosystem differ at each stage of invasion. Journal of Ecology 97, 657–65.

Dehnen-Schmutz K., Touza J., Perrings Perrings. & Williamson M. (2007) The horticultural trade and ornamental plant invasions in Britain. Conservation Biology 21, 224–31.

Díaz S., Kattge J., Cornelissen J. H. C., Wright I. J., Lavorel S., Dray S.…, & Gorné L.D. (2016) The global spectrum of plant form and function. Nature 529, 167–71.

Diez J. M., Williams P. A., Randall R. P., Sullivan J. J., Hulme P.E. & Duncan R. P. (2009) Learning from failures: testing broad taxonomic hypotheses about plant naturalization. Ecology Letters 12, 1174–83.

Dodd A. J., Burgman M. A., McCarthy M.A. & Ainsworth N. (2015) The changing patterns of plant naturalization in Australia. Diversity and Distributions 21, 1038–50.

Drew J., Anderson Anderson. & Andow D. (2010) Conundrums of a complex vector for invasive species control: a detailed examination of the horticultural industry. Biological Invasions 12, 2837–51.

Driscoll D. A., Catford J. A., Barney J. N., Hulme P. E., Inderjit Martin T. G., Pauchard A., Pyšek P., Richardson D. M., Riley Riley. & Visser V. (2014) New pasture plants intensify invasive species risk. Proceedings of the National Academy of Sciences 111, 16622–7.

Duncan R. P. (2011) Propagule pressure. In: Encyclopedia of biological invasions (eds D. Simberloff and M. Rejmanek) Vol. 3 pp. 561–3. University of California Press.

Engemann K., Sandel B., Boyle B., Enquist B. J., Jørgensen P. M., Kattge J., McGill B. J., Morueta-Holme N., Peet R. K., Spencer N. J., Violle C., Wiser S.K. & Svenning J.-C. (2016) A plant growth form dataset for the New World. Ecology 97, 3243.

Essl F., Bacher S., Blackburn T. M., Booy O., Brundu G., Brunel S., Cardoso A.-C., Eschen R., Gallardo B., Galil B., García-Berthou E., Genovesi P., Groom Q., Harrower C., Hulme P. E., Katsanevakis S., Kenis M., Kühn I., Kumschick S., Martinou A. F., Nentwig W., O’Flynn C., Pagad S., Pergl J., Pyšek P., Rabitsch W., Richardson D. M., Roques A., Roy H. E., Scalera R., Schindler S., Seebens H., Vanderhoeven S., Vilà M., Wilson J. R. U., Zenetos Zenetos. & Jeschke J. M. (2015) Crossing frontiers in tackling pathways of biological invasions. BioScience 65, 769–82.

Essl F., Moser D., Dullinger S., Mang Mang. & Hulme P. E. (2010) Selection for commercial forestry determines global patterns of alien conifer invasions. Diversity and Distributions 16, 911–21.

FitzJohn R. G., Pennell M. W., Zanne A. E., Stevens P. F., Tank D.C. & Cornwell W. K. (2014) How much of the world is woody? Journal of Ecology 102, 1266–72.

Funk J. L. & Throop H. L. (2010) Enemy release and plant invasion: patterns of defensive traits and leaf damage in Hawaii. Oecologia 162, 815–23.

Gallagher R. V., Randall R. P. & Leishman M. R. (2015) Trait differences between naturalized and invasive plant species independent of residence time and phylogeny. Conservation Biology 29, 360–9.

Godoy O., Valladares Valladares. & Castro-Díez P. (2012) The relative importance for plant invasiveness of trait means, and their plasticity and integration in a multivariate framework. New Phytologist 195, 912–22.

Goodwin B. J.,McAllister A.J. & Fahrig L. (1999) Predicting invasiveness of plant species based on biological information. Conservation Biology 13, 422–6.

Hamilton M., Murray B., Cadotte M., Hose G., Baker A., Harris Harris. & Licari D. (2005) Life-history correlates of plant invasiveness at regional and continental scales. Ecology letters 8, 1066–74.

Hanspach J., Kühn I., Pyšek P., Boos Boos. & Klotz S. (2008) Correlates of naturalization and occupancy of introduced ornamentals in Germany. Perspectives in Plant Ecology, Evolution and Systematics 10, 241–50.

Heger Heger. & Trepl L. (2003) Predicting Biological Invasions. Biological Invasions 5, 313–21.

Hulme P. E., Bacher S., Kenis M., Klotz S., Kühn I., Minchin D., Nentwig W., Olenin S., Panov V., Pergl J., Pyšek P., Roques A., Sol D., Solarz Solarz. & Vilà M. (2008) Grasping at the routes of biological invasions: a framework for integrating pathways into policy. Journal of Applied Ecology 45, 403–14.

Jeschke J., Aparicio L. G., Haider S., Heger T., Lortie C., Pyšek P. & Strayer D. (2012) Support for major hypotheses in invasion biology is uneven and declining. NeoBiota 14, 1–20.

Kattge J., Diaz S., Lavorel S., Prentice I. C., Leadley P., Bonisch G.…, & Wirth C. (2011) TRY – a global database of plant traits. Global Change Biology 17, 2905–35.

Kendal D., Williams K.J.H. & Williams N. S. G. (2012) Plant traits link people’s plant preferences to the composition of their gardens. Landscape and Urban Planning 105, 34–42.

Kitajima K., Fox A. M., Sato Sato. & Nagamatsu D. (2006) Cultivar selection prior to introduction may increase invasiveness: evidence from Ardisia crenata. Biological Invasions 8, 1471–82.

Klinerová T., Tasevová K. & Dostál P. (2018) Large generative and vegetative reproduction independently increases global success of perennial plants from Central Europe. Journal of Biogeography 45, 1550–9.

Knapp Knapp. & Kühn I. (2012) Origin matters: widely distributed native and non-native species benefit from different functional traits. Ecology Letters 15, 696–703.

Kowarik Kowarik. & von der Lippe M. (2007) Pathways in plant invasions. In: Biological Invasions (ed W. Nentwig) pp. 29–47. Springer.

Kueffer C., Pyšek P. & Richardson D. M. (2013) Integrative invasion science: model systems, multi-site studies, focused meta-analysis and invasion syndromes. New Phytologist 200, 615–33.

Lavoie C., Joly S., Bergeron A., Guay Guay. & Groeneveld E. (2016) Explaining naturalization and invasiveness: new insights from historical ornamental plant catalogs. Ecology and Evolution 6, 7188–98.

Leishman M. R., Wright I. J., Moles A.T. & Westoby M. (2000) The evolutionary ecology of seed size.In: Seeds: the ecology of regeneration in plant communities. 2nd edition (ed M. Fenner) pp. 31-57. CABI Publishing.

Lockwood J. L., Cassey Cassey. & Blackburn T. M. (2005) The role of propagule pressure in explaining species invasions. Trends in Ecology & Evolution 20, 223–8.

Lockwood J. L., Cassey Cassey. & Blackburn T. M. (2009) The more you introduce the more you get: the role of colonization pressure and propagule pressure in invasion ecology. Diversity and Distributions 15, 904–10.

Lockwood J. L., Hoopes M.F. & Marchetti M. P. (2013) An introdution to invasion ecology. In: Invasion Ecology. 2nd edition. (eds J. L. Lockwood, M. F. Hoopes and M. P. Marchetti) pp. 16–44. Wiley-Blackwell, UK.

Lonsdale W. M. (1994) Inviting trouble: Introduced pasture species in northern Australia. Australian Journal of Ecology 19, 345–54.

Martin P. H., Canham C.D. & Marks P. L. (2009) Why forests appear resistant to exotic plant invasions: intentional introductions, stand dynamics, and the role of shade tolerance. Frontiers in Ecology and the Environment 7, 142–9.

Maurel N., Hanspach J., Kühn I., Pyšek P. & van Kleunen M. (2016) Introduction bias affects relationships between the characteristics of ornamental alien plants and their naturalization success. Global Ecology and Biogeography 25, 1500–9.

McGregor K. F., Watt M. S., Hulme P.E. & Duncan R. P. (2012) What determines pine naturalization: species traits, climate suitability or forestry use? Diversity and Distributions 18, 1013–23.

Moles A. T. (2018) Being John Harper: using evolutionary ideas to improve understanding of global patterns in plant traits. Journal of Ecology 106, 1–18.

Moles A. T., Ackerly D. D., Tweddle J. C., Dickie J. B., Smith R., Leishman M. R., Mayfield M. M., Pitman A., Wood J.T. & Westoby M. (2007) Global patterns in seed size. Global Ecology and Biogeography 16, 109–16.

Moravcová L., Pyšek P., Jarošík V., Havlíčková V. & Zákravský P. (2010) Reproductive characteristics of neophytes in the Czech Republic: traits of invasive and noninvasive species. Preslia 82, 365–90.

Muller-Landau H. C. (2010) The tolerance–fecundity trade-off and the maintenance of diversity in seed size. Proceedings of the National Academy of Sciences 107, 4242–7.

Mulvaney M. (2001) The effect of introduction pressure on the naturalization of ornamental woody plants in South-Eastern Australia. In: Weed risk assessment (eds R. H. Groves, F. D. Panetta and J. G. Virtue) pp. 186–93. CSIRO Publishing, Victoria, Australia.

Osone Y., Ishida Ishida. & Tateno M. (2008) Correlation between relative growth rate and specific leaf area requires associations of specific leaf area with nitrogen absorption rate of roots. New Phytologist 179, 417–27.

Pärtel M., Szava-Kovats R. & Zobel M. (2011) Dark diversity: shedding light on absentspecies. Trends in Ecology & Evolution 26, 124–8.

Pergl J., Sádlo J., Petřík P., Danihelka J., Chrtek Jr. J., Hejda M., Moravcová L., Perglová I., Štajerová K. & Pyšek P. (2016a) Dark side of the fence: ornamental plants as a source of wild-growing flora in the Czech Republic. Preslia 88, 163–84.

Pergl J., Pyšek P., Bacher S., Essl F., Genovesi P., Harrower C. A., Hulme P. E., Jeschke J. M., Kenis M., Kühn I., Perglová I., Rabitsch W., Roques A., Roy D. B., Roy H. E., Vilà M., Winter Winter. & Nentwig W. (2017) Troubling travellers: are ecologically harmful alien species associated with particular introduction pathways? NeoBiota 32, 1–20.

Poorter L., Wright S. J., Paz H., Ackerly D. D., Condit R., Ibarra-Manríquez G., Harms K. E., Licona J. C., Martínez-Ramos M., Mazer S. J., Muller-Landau H. C., Peña-Claros M., Webb C.O. & Wright I. J. (2008) Are functional traits good predictors of demographic rates? Evidence from five Neotropical forests. Ecology 89, 1908–20.

Pyšek P., Jarošík V. & Pergl J. (2011) Alien plants introduced by different pathways differ in invasion success: Unintentional introductions as a threat to natural areas. PLOS ONE 6, e24890.

Randall R. P. (2012) A Global Compendium of Weeds. 2nd Edition. Department of Agriculture and Food. Western Australia.

Reich P. B. (2014) The world-wide ‘fast–slow’ plant economics spectrum: a traits manifesto. Journal of Ecology 102, 275–301.

Richardson D. M., Pyšek P. & Carlton J. T. (2011) A compendium of essential concepts and terminology in invasion ecology. In: Fifty Years of Invasion Ecology: The Legacy of Charles Elton (ed D. M. Richardson) pp. 409–20.

Richardson D. M., Pyšek P., Rejmánek M., Barbour M. G., Panetta F.D. & West C.J. (2000) Naturalization and invasion of alien plants: concepts and definitions. Diversity and Distributions 6, 93–107.

Roberts R. E., Clark D.L. & Wilson M. V. (2010) Traits, neighbors, and species performance in prairie restoration. Applied Vegetation Science 13, 270–9.

Schlaepfer D. R., Glättli M., Fischer Fischer. & van Kleunen M. (2010) A multi-species experiment in their native range indicates pre-adaptation of invasive alien plant species. New Phytologist 185, 1087–99.

Simberloff D. (2009) The role of propagule pressure in biological invasions. Annual Review of Ecology, Evolution, and Systematics 40, 81–102.

Šímová I., Violle C., Svenning J.-C., Kattge J., Engemann K., Sandel B., Peet R. K., Wiser S. K., Blonder B., McGill B. J., Boyle B., Morueta-Holme N., Kraft N. J. B., van Bodegom P. M., Gutiérrez A. G., Bahn M., Ozinga W. A., Tószögyová A. & Enquist B. J. (2018) Spatial patterns and climate relationships of major plant traits in the New World differ between woody and herbaceous species. Journal of Biogeography 45, 895–916.

Speek T. A. A., Lotz L. A. P., Ozinga W. A., Tamis W. L. M., Schaminée J.H.J. & van der Putten W. H. (2011) Factors relating to regional and local success of exotic plant species in their new range. Diversity and Distributions 17, 542–51.

Tomasetto F., Duncan R.P. & Hulme P. E. (2013) Environmental gradients shift the direction of the relationship between native and alien plant species richness. Diversity and Distributions 19, 49–59.

Townsley-Brascamp W. & Marr N. E. (1995) Evaluation and analysis of consumer preferences for outdoor ornamental plants. Acta horticulturae 391, 199–206.

van Kleunen M., Dawson Dawson. & Maurel N. (2015) Characteristics of successful alien plants. Molecular Ecology 24, 1954–68.

van Kleunen M., Essl F., Pergl J., Brundu G., Carboni M., Dullinger S., Early R., González-Moreno P., Groom Q. J., Hulme P. E., Kueffer C., Kühn I., Máguas C., Maurel N., Novoa A., Parepa M., Pyšek P., Seebens H., Tanner R., Touza J., Verbrugge L., Weber E., Dawson W., Kreft H., Weigelt P., Winter M., Klonner G., Talluto M.V. & Dehnen-Schmutz K. (2018b) The changing role of ornamental horticulture in alien plant invasions. Biological Reviews 93, 1421–37.

van Kleunen M., Johnson S.D. & Fischer M. (2007) Predicting naturalization of southern African Iridaceae in other regions. Journal of Applied Ecology 44, 594–603.

van Kleunen M., Weber Weber. & Fischer M. (2010b) A meta-analysis of trait differences between invasive and non-invasive plant species. Ecology Letters 13, 235–45.

Westoby M., Falster D. S., Moles A. T., Vesk P.A. & Wright I. J. (2002) Plant ecological strategies: some leading dimensions of variation between species. Annual Review of Ecology and Systematics 33, 125–59.

Wilson J. R. U., Dormontt E. E., Prentis P. J., Lowe A.J. & Richardson D. M. (2009b) Something in the way you move: dispersal pathways affect invasion success. Trends in Ecology & Evolution 24, 136–44.

Zanne A. E., Tank D. C., Cornwell W. K., Eastman J. M., Smith S. A., FitzJohn R. G.…, & Beaulieu J. M. (2014) Three keys to the radiation of angiosperms into freezing environments. Nature 506, 89-92.

Zenni R.D.& Nuñez M. A. (2013) The elephant in the room: the role of failed invasions in understanding invasion biology. Oikos 122, 801–15.

