## Supplementary Material for "Introduction bias: Imbalance in species introductions may obscure the identification of traits associated with invasiveness"

Estibaliz Palma, Jian Yen, Peter A. Vesk, Monserrat Vilà, Jane A. Catford

**Panel A1** Invasions as individual, population or species-level phenomena.

While species introductions are individual-level events, establishment and persistence are population or species-level events (Blackburn et al. 2011). Invasions are commonly understood as species-level phenomena across the scientific and management literature (i.e. a species becomes naturalized, a species becomes invasive). Due to the individual vs. species dichotomy, trait distributions may be drawn using either individual-level values or species-level mean values. This decision may have considerable implications, especially for the introduction stage: an individual-level approach accounts for the strength with which each species (and therefore their trait values) have been introduced, a species-level approach assumes that each species has been introduced equally (species contribute a single trait value to the distribution, i.e. their mean). The former approach seems more appropriate to accurately represent the distribution of trait values that have been introduced by incorporating the propagule pressure for each introduced species. However, information on introduction patterns is usually not easily available, and therefore taking an individual-level approach is usually challenging.

The theoretical framework presented in this paper (Figure 3) assumes that species' probability to be transported and introduced is independent from their distributional range and abundance worldwide; thus, it does not incorporate the idea that the propagule pressure for each introduced species is unequal. To examine the implications of this assumption, we compare the worldwide distribution of four traits - SLA, height, seed mass and woodiness - when species worldwide availability (and therefore the probability that they are picked up and introduced outside their natural range) is either ignored or acknowledged (Figure A1). We concluded that the shape of the trait distribution based on species-level mean values and the shape of the distribution based on weighted mean values were similar enough to assume little effect of the chosen approach on the conclusions drawn from the theoretical framework.

**Panel A2** Code for simulations

### Define probability functions

```
normal_probability <- function (x, mean, sd) {
	out <- exp(( -(x - mean) ** 2) / (2 * (sd ** 2)))
	out <- out / sum(out)
	out
	}
```

```
linear_probability <- function (x, intercept, slope) {
	out <- intercept + slope * x
	out <- out / sum(out)
	out
	}
```

```
log_linear_probability <- function (x, intercept, slope) {
	out <- exp(intercept + slope * x)
	out <- out / sum(out)
	out
	}
```

### Simulation of introduction scenarios

```
Random_introduction <- runif (10000, min = 1, max = 100)
```

```
Biological_bias <- rlnorm (10000, mean_norm =20, sd_norm = 10)
```

```
Human_preference_bias <- sample (x = Biological_bias, size = 10000, prob =
normal_probability (Biological_bias, mean=50, sd=10), replace = TRUE)
```

Human_preference_bias: Mean for main framework: 40
Means for extension of the framework: 30, 45, 60

### Simulation of establishment scenarios

```
Established_pool_noeffect <- sample (x = Introduced_pool, size = 1000, prob = rep
(1, length (Introduced_pool)), replace = FALSE)
```

```
Established_pool_effect <- sample (x = Introduced_pool, size =1000, prob =
linear_probability (Introduced_pool, intercept = 0, slope = 0.005), replace =
FALSE)
```

Introduced_pool can be Random_introduction, Biological_bias or Human_preference_bias
Effect for main framework: 0.005
Effects for extension of the framework:
`prob = log_linear_probability (Introduced_pool, intercept, slope)`
`slope` = 0.0025, 0.05, 0.25

### Simulation of invasion scenarios

```
Invasive_pool_noeffect <- sample (x = Established_pool, size = 100, prob = rep (1,
length (Established_pool)), replace = FALSE)
```

```
Invasive_pool_effect <- sample (x = Established_pool, size = 100, prob =
linear_probability (Established_pool, intercept = 0, slope = 0.005), replace =
FALSE)
```

Effect for main framework: 0.005
Effects for extension of the framework:
`prob = log_linear_probability (Established_pool, intercept, slope)`
`slope` = 0.0025, 0.05, 0.25

**Table A1** Descriptors of distributions presented in the theoretical framework (Figure 3), including mean, standard deviation and kurtosis. Kurtosis of a standard normal distribution is 3. Values of kurtosis above 3 represent a pointier distribution and values below 3, a flatter distribution.

| | | Random introduction | | | Biologically biased introduction | | | Human biased introduction | | |
|---|---|---|---|---|---|---|---|---|---|---|
| | | **Mean** | **SD** | **Kurtosis** | **Mean** | **SD** | **Kurtosis** | **Mean** | **SD** | **Kurtosis** |
| **A.** | **INTRODUCTION** | | | | | | | | | |
| | Introduced pool | 50.476 | 28.614 | 1.797 | 19.992 | 10.264 | 14.678 | 39.254 | 9.742 | 2.987 |
| **B.** | **NATURALIZATION - trait involved** | | | | | | | | | |
| | Established pool | 66.499 | 23.247 | 2.372 | 24.310 | 13.278 | 29.946 | 41.628 | 9.710 | 2.859 |
| | Failures | 48.695 | 28.602 | 1.800 | 19.512 | 9.756 | 6.994 | 38.990 | 9.710 | 3.004 |
| **D.** | **INVASION - trait involved** | | | | | | | | | |
| | Invasive pool | 76.249 | 17.179 | 2.529 | 30.320 | 17.445 | 11.441 | 44.853 | 10.308 | 2.216 |
| | Non-invasive pool | 65.416 | 23.586 | 2.291 | 23.642 | 12.568 | 35.955 | 41.270 | 9.580 | 2.993 |
| **E.** | **INVASION - trait irrelevant** | | | | | | | | | |
| | Invasive pool | 67.049 | 22.492 | 2.214 | 25.398 | 16.507 | 15.596 | 41.487 | 9.229 | 3.097 |
| | Non-invasive pool | 66.438 | 23.341 | 2.384 | 24.189 | 12.875 | 32.934 | 41.644 | 9.766 | 2.834 |
| **C.** | **NATURALIZATION - trait irrelevant** | | | | | | | | | |
| | Established pool | 48.674 | 28.428 | 1.819 | 20.511 | 10.203 | 5.773 | 39.941 | 9.586 | 3.086 |
| | Failures | 50.676 | 28.630 | 1.794 | 19.934 | 10.269 | 15.665 | 39.178 | 9.756 | 2.976 |
| **F.** | **INVASION - trait involved** | | | | | | | | | |
| | Invasive pool | 64.252 | 25.404 | 2.262 | 26.150 | 13.329 | 5.536 | 41.768 | 9.904 | 2.919 |
| | Non-invasive pool | 46.944 | 28.231 | 1.841 | 19.884 | 9.602 | 4.831 | 39.738 | 9.534 | 3.123 |
| **G.** | **INVASION - trait irrelevant** | | | | | | | | | |
| | Invasive pool | 47.008 | 29.605 | 1.796 | 22.630 | 12.780 | 5.830 | 40.955 | 9.383 | 2.899 |
| | Non-invasive pool | 48.860 | 28.305 | 1.825 | 20.275 | 9.856 | 5.334 | 39.828 | 9.606 | 3.097 |

**Figure A1** Distribution of worldwide values of specific leaf area, height and seed mass, and frequency of categories woody and non-woody. Continuous lines and black bars represent the relative frequency of trait values when each species contributes equally to the distributions (one species = one record); dashed lines and white bars represent the relative frequency of trait values when species' contributions are weighted based on species' occurrence worldwide (one species = as many records as number of countries where it has been found, so species with large geographic range will contribute more). The datasets include 2,912 (SLA); 3,456 (height); 2,446 (seed mass) and 5,313 species (woodiness). Records of species' trait and species' global distribution were collected from the BIEN database (Enquist et al. 2016, Maitner et al. 2018).

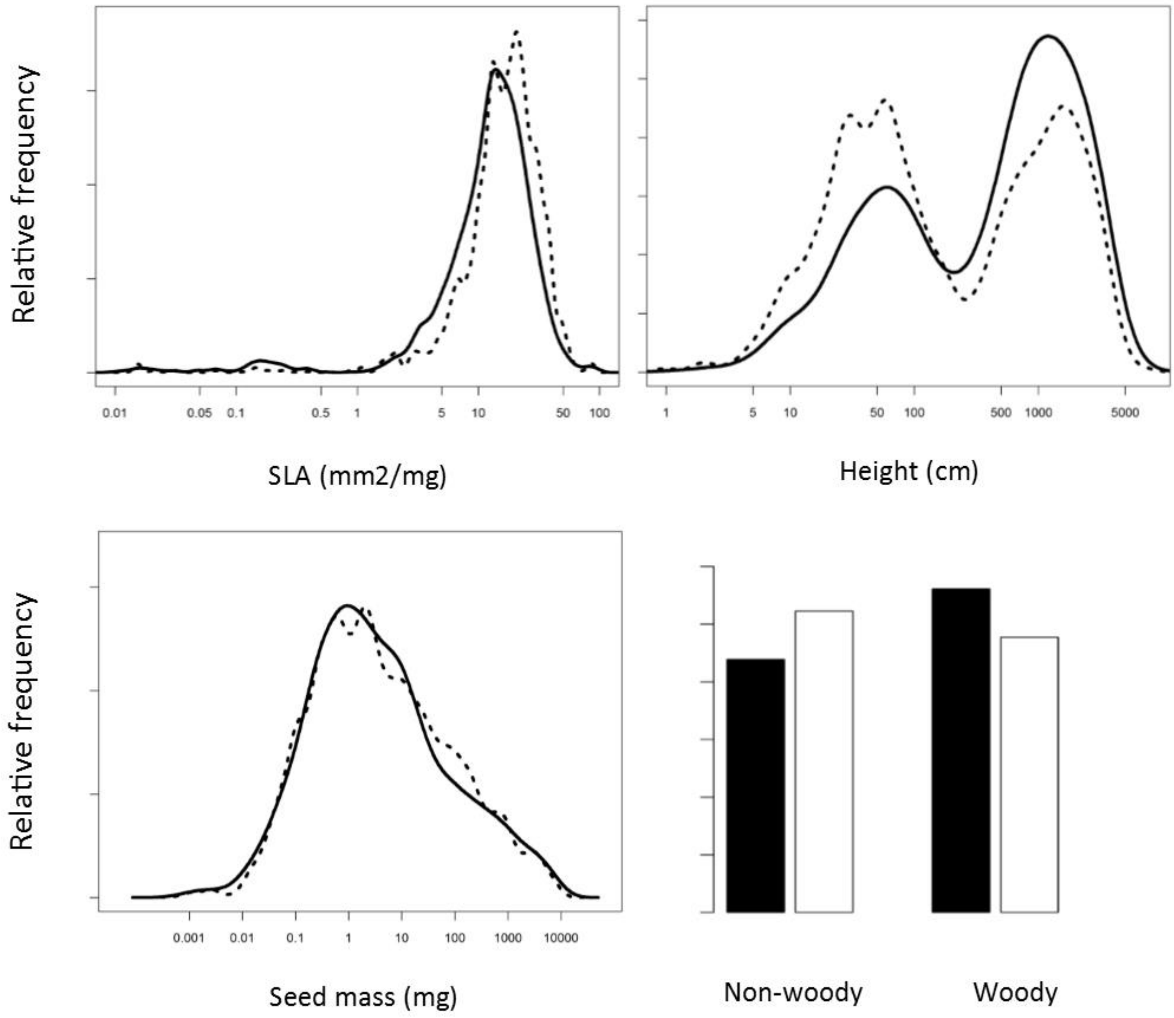
